# The central amygdala integrates exogenous glucagon-like peptide 1 signals

**DOI:** 10.64898/2026.04.06.716705

**Authors:** Miguel Duran, Ningxiang Zeng, Elam J. Cutts, Kirk Habegger, J. Andrew Hardaway

## Abstract

Nuclei within the limbic system like the central amygdala (CeA) play a critical role in mediating fear, motivation, reward, and appetitive behavior. Although previous reports demonstrate the presence of the glucagon-like peptide-1 receptor (GLP-1R) in limbic nuclei, how limbic neurons mediate the actions of systemically administrated GLP-1R agonists is unclear. In this study, we investigated the CeA’s response to peripherally administered GLP-1R agonist Exendin-4 (Ex-4) *in vivo*, and determined the functional requirement of select CeA neuron populations in acute Ex-4 induced hypophagia. Using fiber photometry, we observed that Ex-4 promoted a rapid and lasting activation of CeA neurons that was blocked by pretreatment with the GLP-1R antagonist Exendin-9. We then tested the functional requirement of CeA neuron activation in mediating Ex-4 induced hypophagia of standard grain chow using inhibitory chemogenetics. Chemogenetic inhibition of all CeA neurons significantly suppressed the hypophagic actions of Ex-4. Then using selective mouse Cre-drivers, we found that chemogenetic inhibition of protein kinase c delta (*Prk-cd* ^CeA^) and GLP-1R (*Glp1r* ^CeA^), but not somatostatin (*Sst*^CeA^), neurons also attenuates the full hypophagic effect of Ex-4. Having observed that inhibition of *Glp1r*^CeA^ modestly attenuated Ex-4 induced hypophagia of standard chow, we then tested whether these neurons might mediate Ex-4 suppression of energy-dense, palatable diet. We used intermittent high-fat diet (HFD) access and found that inhibition of *Glp1r*^CeA^ neurons significantly rescued the reduction of HFD consumption by Ex-4. Collectively, these data demonstrate that the CeA responds to peripherally administered GLP-1R agonists and that multiple CeA neuron mediate GLP-1R agonist-mediated hypophagia.

## 1. Introduction

Type 2 diabetes and obesity are serious public health maladies that affect many developed nations. In the U.S., 25% of the population are obese [1], which brings increased risk for more serious health complications, e.g., cardiovascular disease, cancer, and the development of type 2 diabetes (National Heart, Lung, and Blood Institute 2013). Over the past ten years, the FDA has approved glucagon-like peptide-1 receptor (GLP-1R) agonists like liraglutide or semaglutide for the treatment of type 2 diabetes and weight management [2]. These drugs promote insulin release, delay gastric emptying, and decrease appetite, the latter of which is mediated through the central nervous system (CNS) [3], [4], [5], [6], [7]. However, the precise circuitry and mechanisms in the CNS recruited by GLP-1R agonists to reduce food intake have yet to be fully characterized. In particular, further research is needed to establish the direct vs indirect recruitment of neural circuits by GLP-1R agonists. Several groups have characterized the expression and action of GLP-1R-expressing cells within the CNS and their effects on physiology and behavior [3], [4], [5], [6], [7], [7], [8], [9], [10], [11], [12], [13], [14], [15], [16], [17], [18], [19], [20],[21], [22], [23], [24], [25], [26], [27], [28], [29], [30], [31], [32], [33], [34], [35], [36], [37], [38], [39], [40], [41], [42], [43], [44], [45], [46], [47], [48], [49], [50], [51], [52], [53], [54], [55], [56], [57]. GLP-1R-expressing cells are evolutionarily conserved and enriched in brain nuclei that respond to exogenous GLP-1R agonists [8], [9], [10], [30], [37], [58], [59], [60], [61], [62]. Multiple studies demonstrate that the hypothalamus and hind-brain contain many GLP-1R+ cells and that these nuclei are required to mediate the anorexigenic effects of GLP-1R agonists [3], [5], [20], [21], [26], [58]. However, GLP-1R expression is also enriched in limbic sites such as the bed nucleus of the stria terminalis (BNST), lateral septum (LS), and central nucleus of the amygdala (CeA) which are key nuclei that regulate motivation, reward, and appetitive behavior [15], [39], [48], [53], [63]. While GLP-1R-expressing or other cells within the BNST, LS, and CeA regulate feeding, the role of neuronal populations in these limbic nuclei in mediating responses to exogenous GLP-1R agonists is less clear [39], [48], [53], [57].

The CeA is a heterogeneous nucleus containing various GABAergic cell types that express distinct signaling molecules and neuropeptides [64], [65], [66], [67], [68], [69], [70], [71], [72]. The CeA’s composition and input/output architecture allows it to function as a hub that integrates local signaling while also sending and receiving projections throughout the CNS [73], [74]. Our lab has shown that CeA GLP-1R-expressing neurons are distinct from other known genetically-defined neurons in the CeA, are enriched within the medial division of the CeA, and have unique electrophysiological properties [54]. Of particular note, multiple Fos studies have shown that, regardless of the route of administration, the type or dose of GLP-1R agonist used, species, or subject sex, neurons in the CeA are consistently activated [75]. Despite these reports, the CeA has not received more attention in the effort to understand the brain mechanisms mediating hypophagia by exogenously administered GLP-1R agonists. In this study, we tested whether distinct, genetically-defined neuronal populations within the CeA are required for the hypophagic effect of peripheral GLP-1R agonist administration.

## 2. Results

### 2.1 Peripheral GLP-1R signaling activates CeA neurons, in vivo

Previous reports using *post hoc* Fos immunolabeling demonstrate that peripheral and intracerebroventricular administration of GLP-1R agonists consistently activate the CeA [75]. To investigate this activation *in vivo*, we used the genetically-encoded calcium indicator GCaMP7 paired with fiber photometry recordings to capture the CeA’s response to systemic administration of an GLP-1R agonist, Exendin-4 (Ex-4) in freely behaving mice. The CeA is primarily a GABAergic nucleus, therefore we bilaterally injected *Slc32a1(vGat)*-IRES-Cre mice with AAV-hSyn-FLEX-GCaMP7f into the CeA followed by bilateral implantation of fiber optics dorsal to the injection site (**Figure 1A**), resulting in the expression of GCaMP7f throughout the CeA and correctly targeted fiber optics (**Figure 1B**). Following recovery from surgery, habituation to behavior chambers, and handling, mice were subjected to randomized treatment (i.p.) of either Saline and Saline (Sal/Sal), Saline and Ex-4 (5 µg/kg) (Sal/Ex-4), Exendin-9 (50 µg/kg) and Saline (Ex-9/Sal), and Ex-9 and Ex-4 (Ex-9/Ex-4) during recordings. As expected due to the CeA’s role in encoding pain and aversive stimuli, all mice showed strong and rapid responses to the injection itself (**Figure 1C**). For Sal/Ex-4 treatment; however, we observed a rapid and sustained activation of CeA neurons following injection of Ex-4 when compared to Sal/Sal, Ex-9/Ex-4, and Ex-9/Sal (Figure 1C, Ex-9/Sal not shown). Examination of Ex-4 responses in individual mice revealed that a majority of mice show sustained and prominent when compared to Saline alone, but some mice did not show responses (**Figure 1D**). Quantification of the net AUC from 15-30’ post-injection of the 2nd treatment, where the animal’s response to injection has subsided, we observed a significant increase in the Sa/ Ex-4 treatment condition when compared to Sal/Sal, as well as a non-significant increase when compared to Ex-9/Ex-4 and Ex-9/Sal (**Figure 1E**). We observed no significant difference in the peak response between any of the treatments (**Figure 1F**). In *post hoc* examination of the CeA in these mice, we confirmed GCaMP7 expression and documented fiber optic termination sites (Figure 1G-J). Collectively, these data demonstrate that systemic administration of the GLP-1R agonist Exendin-4 results in sustained activation of the CeA.

**Figure 1.**
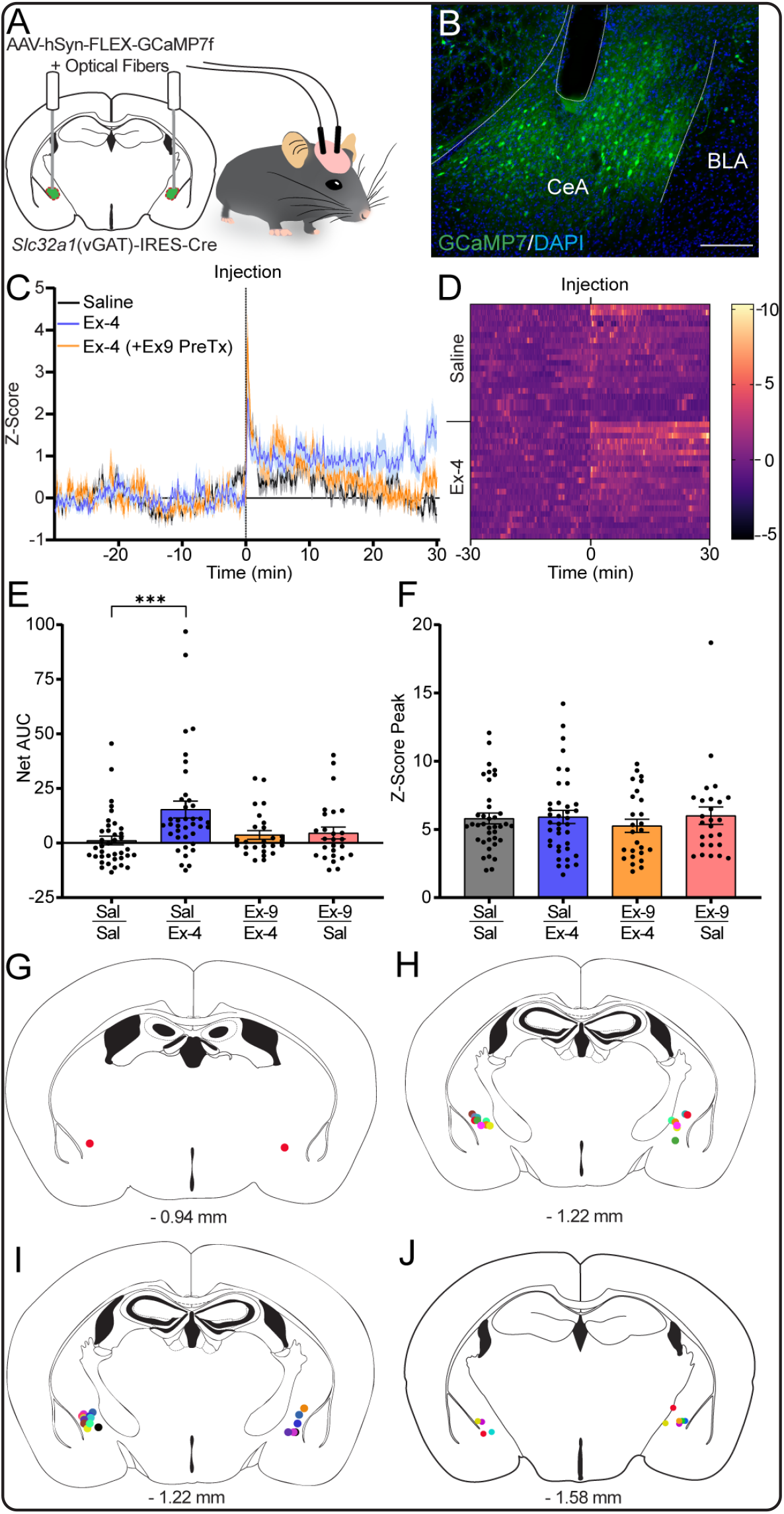
Systemic administration of the long acting GLP-1R agonist Exendin-4 results in lasting activation of CeA neurons in vivo. **A.**Surgical schematic of bilateral injection of AAV-hSyn-FLEX-GCaMP7 and fiber optic implants in CeA of Sc32a1(*vGat*)-Cre mice (n = 15-21). **B**. Representative image of GCaMP7 expression in the CeA and fiber optic termination site from a *vGat*-Cre injected mouse taken at 20x magnification. Scale bar = 200 µM. **C**. Z-Score traces of GCaMP7 fluorescence during a 1 hr window surrounding each respective treatment (i.p.). **D**. Heatmap of calcium transients during 1Hr window for each animal during Saline or Ex-4 treatment (n = 21). Data were sorted by descending area-under-the-curve calculations. **E**. Quantification of the net area under the curve (AUC) 15-30 minutes post-injection for each treatment (n = 15-21). p = 0.0012; Post-test: Sal/Sal vs Sal/Ex4: p = 0.0006; Sal/Sal vs Sal/Ex9: p > 0.9999; Sal/Sal vs Ex-9/Ex4: p > 0.9999 **F**. Quantification of the maximum Z-Score peak post-injection for each treatment (n = 15-21). p = 0.7919. **G-J**. Schematic representations of fiber optic termination sites at the CeA for each mouse located at -0.94 (G), -1.22 (H-I), and -1.58 (J) mm for mouse brain. For E-F, data were analyzed using Kruskal-Wallis test with Dunn’s multiple comparisons. *p < 0.05, **p < 0.01, ***p < 0.001

### 2.2 Inhibition of *vGat*^CeA^ neurons attenuates GLP-1-induced hypophagia

Having observed a rapid and sustained activation of CeA neurons in response to a systemic GLP-1R agonist, we tested the requirement of CeA neuron activation in Ex-4-mediated hypophagia. We used a chemogenetic strategy to reversibly silence all CeA neurons prior to Ex-4 administration. As the CeA is a GABAergic nucleus, we bilaterally injected *vGat*-Cre mice with either AAV-hSyn-DIO-mCherry or AAV-hSyn-DIO-hM4d-mCherry into the CeA (**Figure 2A**), resulting in mCherry expression throughout the CeA (**Figure 2B**). To validate the function of hM4d, we used whole-cell patch clamp electrophysiology in *ex vivo* CeA brain slices and recorded from hM4d-mCherry+ neurons (**Figure 2C**). In current clamp mode, bath application of the designer ligand deschloroclozapine (DCZ, 1 µM) resulted in significant hyperpolarization after a 5-minute baseline recording (**Figure 2D**). These data demonstrate that activation of hM4D by DCZ results in inhibition of neural activity and silencing of CeA neurons.

**Figure 2.**
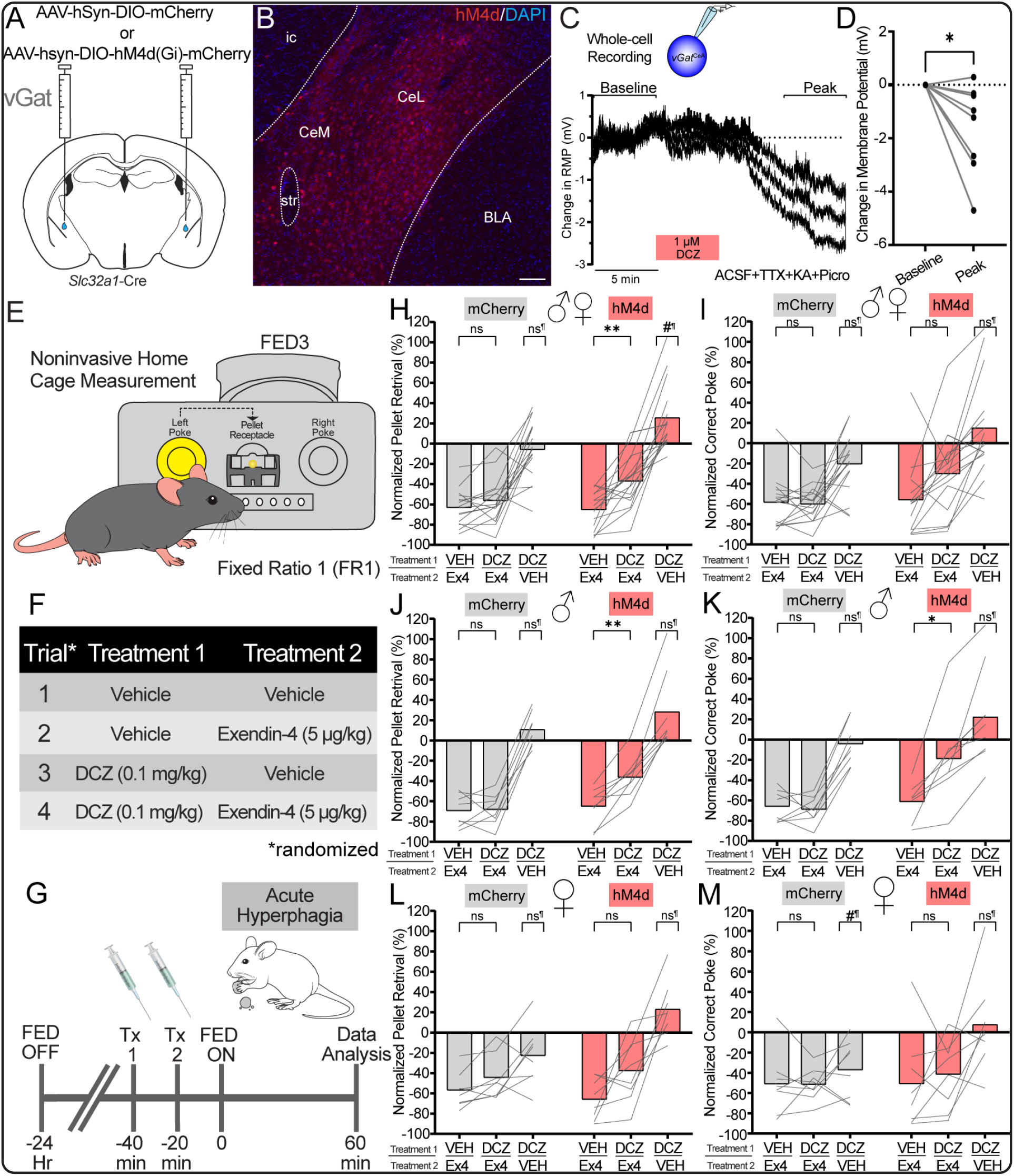
Chemogenetic inhibition of CeA neurons attenuates systemic Exendin-4-mediated hypophagia. **A.**Surgical schematic of bilateral injection of either AAV-hSyn-DIO-mCherry (n = 8/sex) or AAV-hSyn-hM4d-mCherry (n = 8/sex) in the CeA of Sc32a1(vGat)-Cre mice. **B**. Representative image of hM4d-mCherry expressing CeA tissue from *vGat*-Cre injected mouse taken at 20x magnification. Scale bar = 200 µM. **C**. Average electrophysiological trace of vGatCeA neurons in current clamp using whole-cell patch-clamp following DCZ (1 µM) wash-on in the presence of tetrodotoxin and synaptic blockers. **D**. Quantification of change in membrane potential (C) pre- and post-DCZ wash-on. n = 8 cells. Wilcoxon test *p* = 0.0156. **E**. Representative illustration of mouse interacting with Feeding Experimental Device (FED3) under fixed-ratio 1 paradigm in order to receive a food pellet. **F**. Table of treatment (Tx) regime that is randomized for each animal. **G**. Timeline for each fasted-refeed/treatment session (i.p.) and endpoint (60 min) for data acquisition for analysis. Each animal is exposed to each Tx regime (Fig. 2F) at least once over the course of 4 sessions. **H**. Pellets retrieved by all *vGat*-Cre mice during each Tx session. Group (Gr, mCherry/hM4d) X Tx: F(2.173, 65.20) = 5.194, *p* = 0.0067; Gr effect: F(1, 30) = 7.226, *p* = 0.0116; Tx effect: F(2.173, 65.20) = 102.2, *p* < 0.0001. Post tests - Veh/Ex4 vs DCZ/ Ex4: mCherry (*p* = 0.3755), hM4 (*p* = 0.0064); DCZ/Veh vs Veh/Veh: mCh (*p* = 0.8494), hM4 (*p* = 0.0294). **I**. Active port pokes by all *vGat*-Cre mice during each Tx session. Gr X Tx: F(2.721, 81.62) = 4.528, *p* = 0.007; Gr effect: F(1, 30) = 5.558, *p* = 0.0251; Tx effect: F(2.721, 81.62) = 45.38, *p* < 0.0001. Post tests - Veh/ Ex4 vs DCZ/Ex4: mCherry (*p* = 0.9862), hM4d (*p* = 0.1180); DCZ/Veh vs Veh/Veh: mCh (*p* = 0.0679), hM4 (*p* = 0.5978). **J**. Pellets retrieved by male *vGat*-Cre mice during each Tx session Gr X Tx: F(1.726, 24.17) = 2.909, *p* = 0.0804; Gr effect: F(1, 14) = 4.386, *p* = 0.0549; Tx effect: F(1.726, 24.17) = 96.91, *p* < 0.0001. Post tests - Veh/Ex4 vs DCZ/Ex4: mCh (*p* = 0.9974), hM4 (*p* = 0.0064); DCZ/Veh vs Veh/Veh: mCh (*p* = 0.3424), hM4 (*p* = 0.2469). **K**. Active port pokes by male *vGat*-Cre mice during each Tx session. Gr X Tx: F(1.838, 25.73) = 4.791, *p* = 0.0192; Gr effect: F(1, 14) = 4.210, *p* = 0.0594; Tx effect: F(1.838, 25.73) = 43.70, *p* < 0.0001. Post tests: Veh/Ex4 vs DCZ/Ex4: mCh (*p* = 0.9594), hM4 (*p* = 0.0382); DCZ/Veh vs Veh/Veh: mCh (*p* =0.9480), hM4 (*p* = 0.6168). **L**. Pellets retrieved by female *vGat*-Cre mice during each Tx session. Gr X Tx: F(2.283, 31.96) = 5.795, *p* = 0.0054; Gr effect: F(1, 14) = 2.559, *p* = 0.1320; Tx effect: F(2.283, 31.96) = 37.68, *p* < 0.0001. Post tests: Veh/Ex4 vs DCZ/Ex4: mCh (*p* = 0.1649), hM4 (*p* = 0.2600); DCZ/Veh vs Veh/Veh: mCh (*p* = 0.2246), hM4 (*p* = 0.1749). **M**. Active port pokes by female *vGat*-Cre mice during each Tx session. Gr X Tx: F(2.017, 28.24) = 2.858, *p* = 0.0736; Gr effect: F(1, 14) = 1.540, *p* = 0.2350; Tx effect: F(2.017, 28.24) = 15.51, *p* < 0.0001. Post tests: Veh/Ex4 vs DCZ/Ex4: mCh (*p* >0.9999), hM4 (*p* = 0.9399); DCZ/Veh vs Veh/Veh: mCh (*p* = 0.0366), hM4 (*p* = 0.9643). For H-M, data were analyzed using two-way repeated measures ANOVA with Tukey’s multiple comparisons. Datapoints for each mouse during each Tx session were normalized to their respective Vehicle/Vehicle session.¶ Indicates a post test to Veh/Veh condition. Raw data shown in Figure S1.

We then investigated the effect of chemogenetic inhibition of the CeA on GLP-1R-agonist-mediated hypophagia. To do this, we used a noninvasive method that allowed quantal assessment of both food intake and food seeking using the open-source FED3 device (**Figure 2E**) [76]. The use of FED3 enabled us to precisely quantify the number of nose pokes and pellet retrievals after delivery of GLP-1R agonists or any drug treatment in individual mice. After surgery, mice were singly-housed and trained to self-feed under a fixed-ratio 1 (FR1) paradigm where one poke in the active port resulted in the delivery of one 20 mg precision grain pellet. We allowed all mice at least one week of self-feeding before beginning drug experiments. To test our hypothesis, we used four randomized drug treatment combinations (**Figure 2F**) in both mCherry (control) and hM4d-mCherry-expressing animals. The dosage of Ex-4 (5 µg/ kg) was titrated to produce a ∼50% suppression in food intake, which allows for modulation of food intake in response to DCZ in either direction. To promote overall food intake, we food-deprived animals for 24 hours prior to an experimental day. On the experimental day, animals received a randomized two-drug treatment regimen prior to FED3 access and refeeding (**Figure 2G**). As expected, we observed that mice consume 30-40 precision pellets and ∼50 active port pokes in the first hour after food deprivation (**Figure 2H-I, S1A-B**), and Ex-4 pretreatment resulted in a 50-60% reduction in both pellets retrieved and active pokes performed in both sexes of mice. We focused on the first hour due to the rapid pharmacokinetics of DCZ [77].

Here we present data in normalized and raw pellet formats in the main figures and supplements, respectively. In control animals, DCZ treatment prior to Ex-4 resulted in no change in pellet retrieval and active pokes relative to Veh/Ex-4 conditions. In hM4d animals; however, we observed that DCZ treatment mitigated the hypophagic effect of Ex-4, but not completely. This resulted in a significant increase of pellets retrieval and active pokes in DCZ/Ex-4 conditions relative to Veh/Ex-4 in the hM4d group. We observed this phenomenon in male hM4d-expressing mice (**Figure 2J-K, S1C-D**), and although a similar trend was apparent in female mice, we failed to detect a significant attenuation of Ex-4 hypophagia in response to DCZ (**Figure 2L-M, S1E-F**). In parallel, we also tested the effect of DCZ alone (DCZ/Veh). Inhibition of CeA neurons produced a subtle and significant increase in pellet retrievals, but not active pokes (**Figure 2H-I, S1A-B**) relative to Veh/Veh, with a nonsignificant increase observed in both male and female hM4d mice (**Figure 2J-M, S1C-F**). Collectively, these data demonstrate that activation of CeA neurons is required for the complete hypophagic effect of the GLP-1R agonist Ex-4.

### 2.3 Inhibition of *Pkcd*^CeA^ neurons rescues GLP-1R agonist-induced hypophagia

The CeA is known for its heterogeneous neuron populations that express different signaling molecules or neuropeptides. Of these neuronal populations, protein kinase C-delta (*Prkcd*) is an enriched gene that marks neurons in the lateral CeA (CeL). Also, *Prkcd*^CeA^ neurons mediate the influence of other anorexigenic signals like cholecystokinin and reduce feeding when activated [78]. With that, we investigated the role of *Prkcd*^CeA^ neurons in mediating GLP-1R agonist-induced hypophagia, specifically whether inhibiting them would attenuate the effect of Ex-4. Similar to our previous strategy to silence all GABAergic neurons in the CeA, we bilaterally injected either mCherry or hM4d into the CeA of *Prkcd*-Cre mice (**Figure 3A**), resulting in mCherry expression in the CeL (**Figure 3B**). Following the same experimental approach (**Figure 2E-G**), we investigated the effects of *Pkcd*^CeA^ neuron inhibition on GLP-1R agonist-mediated hypophagia. In control animals, DCZ treatment prior to Ex-4 resulted in no change in pellets retrieved and active pokes relative to Veh/Ex-4 conditions (**Figure 3C-D, S2A-B**). In hM4d animals; however, there was a significant increase in pellet retrieval and active pokes relative to Veh/ Ex4. Similar to the inhibition of all CeA neurons, attenuation of Ex-4-mediated hypophagia was incomplete. When separated by sex, we observed a significant increase of pellets retrieval in both male and female hM4d mice (**Figure 3E&G, S2C&E**). For active pokes, we only observed a significant increase in female, but not male, hM4d mice (**Figure 3F+H, S2D + F**). When examining the effects of DCZ/Veh alone, inhibition of *Pkcd*^CeA^ neurons significantly increased pellet retrieval and active pokes in hM4d mice (**Figure 3C-D, S2A-B**) relative to Veh/Veh. In male mice, a nonsignificant increase was observed (**Figure 3E-F, S2C-D**), but female mice did significantly increase their pellet retrieval and active pokes (**Figure 3G-H, S2E-F**). Collectively, these data demonstrate that activation of *Prkcd*^CeA^ neurons is required for the complete hypophagic effect of Ex-4.

**Figure 3.**
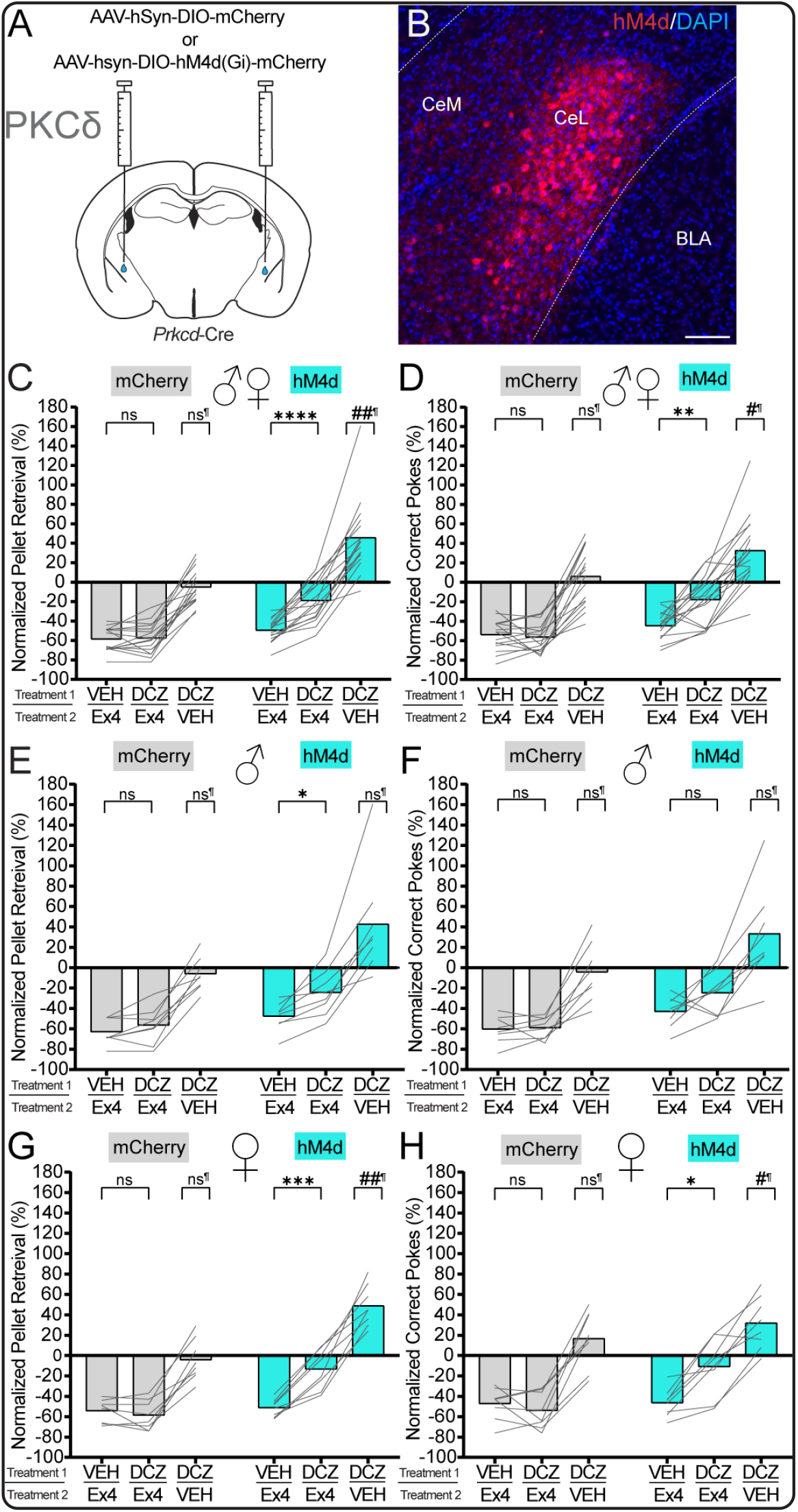
Chemogenetic inhibition of *Prkcd*^CeA^ neurons potently attenuates systemic Exendin-4-mediated hypophagia. **A.**Surgical schematic of bilateral injection of either AAV-hSyn-DIO-mCherry (n = 8/sex) or AAV-hSyn-hM4d-mCherry (n = 8/sex) in the CeA of *Prkcd*-Cre mice. **B**. Representative image of hM4d-mCherry-expressing CeA tissue from Prkcd-Cre mouse taken at 20x magnification. Scale Bar = 200 µM. **C**. Pellets retrieved by all *Prkcd*-Cre mice during each treatment (Tx) session. Group (Gr, mCherry/hM4d) X Tx: F(1.702, 51.06) = 18.03, *p* < 0.0001; Gr effect: F(1, 30) = 31.68, *p* < 0.0001; Tx effect: F(1.702, 51.06) = 145.8, *p* < 0.0001. Post tests: Veh/Ex4 vs DCZ/ Ex4: mCh (*p* = 0.9740), hM4 (*p* < 0.0001); DCZ/Veh vs Veh/Veh: mCh (*p* = 0.6740), hM4 (*p* = 0.0013) **D**. Active port pokes by all *Prkcd*-Cre mice during each Tx session. Gr X Tx: F(2.003, 60.09) = 7.274, *p* = 0.0015; Gr effect: F(1, 30) = 14.91, *p* = 0.0006; Tx effect: F(2.003, 60.09) = 99.40, *p* < 0.0001. Post tests: Veh/Ex4 vs DCZ/Ex4: mCh (*p* = 0.9050), hM4 (*p* = 0.0047); DCZ/Veh vs Veh/Veh: mCh (*p* = 0.8286), hM4 (*p* = 0.0118) **E**. Pellets retrieved by male *Prkcd*-Cre mice during each Tx session. Gr X Tx: F(1.426, 19.96) = 4.670, *p* = 0.0316; Gr effect: F(1, 14) = 9.769, *p* = 0.0074; Tx effect: F(1.426, 19.96) = 49.28, *p* < 0.0001. Post tests: Veh/Ex4 vs DCZ/Ex4: mCh (*p* = 0.3768), hM4 (*p* = 0.0200); DCZ/Veh vs Veh/Veh: mCh (*p* = 0.7379), hM4 (*p* = 0.1827) **F**. Active port pokes by male *Prkcd*-Cre mice during each Tx session. Gr X Tx: F(1.474, 20.64) = 2.889, *p* = 0.0911; Gr effect: F(1, 14) = 10.99, *p* = 0.0051; Tx effect: F(1.474, 20.64) = 39.49, *p* < 0.0001. Post tests: Veh/Ex4 vs DCZ/ Ex4: mCh (*p* = 0.9868), hM4 (*p* = 0.3144); DCZ/Veh vs Veh/Veh: mCh (*p* = 0.9710), hM4 (*p* = 0.2576) **G**. Pellets retrieved by female *Prkcd*-Cre mice during each Tx session. Gr X Tx: F(2.088, 29.24) = 20.65, *p* < 0.0001; Gr effect: F(1, 14) = 29.14, *p* < 0.0001; Tx effect: F(2.088, 29.24) = 125.3, *p* < 0.0001. Post tests: Veh/Ex4 vs DCZ/Ex4: mCh (*p* = 0.6484), hM4 (*p* = 0.0004); DCZ/Veh vs Veh/Veh: mCh (*p* = 0.9369), hM4 (*p* = 0.0010). **H**. Active port pokes by female *Prkcd*-Cre mice during each Tx session. Gr X Tx: F(2.399, 33.58) = 6.529, *p* = 0.0025; Gr effect: F(1, 14) = 4.738, *p* = 0.0471; Tx effect: F(2.399, 33.58) = 66.73, *p* < 0.0001. Post tests: Veh/Ex4 vs DCZ/Ex4: mCh (*p* = 0.7470), hM4 (*p* = 0.0151); DCZ/Veh vs Veh/Veh: mCh (*p* = 0.3738), hM4 (*p* = 0.0372). For C-H, data were analyzed using two-way repeated measures ANOVA with Tukey’s multiple comparisons. Datapoints for each mouse during each treatment session were normalized to their respective Veh/Veh session. ¶ Indicates a post test to Veh/Veh condition. Raw data in Figure S2.

### 2.4 Inhibition of *Sst*^CeA^ neurons has no effect on GLP-1R agonist-induced hypophagia

Somatostatin marks a neuronal population within the CeL that has minimal overlap with *Prkcd*^CeA^ [78]. In mice, *Sst*^CeA^ neurons mediate both appetitive and aversive behaviors and are interconnected with *Pkcd*^CeA^ neurons [68], [79], [80]. Similar to our previous experiments, we targeted *Sst*^CeA^ neurons to determine if they mediate GLP-1R agonist-induced hypophagia. We bilaterally injected either mCherry or hM4d into the CeA of *Sst*-Cre mice (**Figure 4A**), resulting in mCherry expression throughout the CeA, predominantly within the CeL (**Figure 4B**). In both control and hM4d animals, DCZ treatment prior to Ex-4 resulted in no significant differences in either pellet retrieval or active pokes relative to Veh/Ex-4 conditions (**Figure 4C-D, S3A-B**). When separated by sex, we observed no significant differences in male (**Figure 4E-F, S3C-D**) and female (**Figure 4G-H, S3E-F**) mice. When examining the effects of inhibition alone, inhibition of *Sst*^CeA^ neurons significantly decreased pellets retrieved in hM4d male mice of their normalized data (**Figure 4E, DCZ/Veh**) but not in their raw data (**S3C**) relative to Veh/Veh.

**Figure 4.**
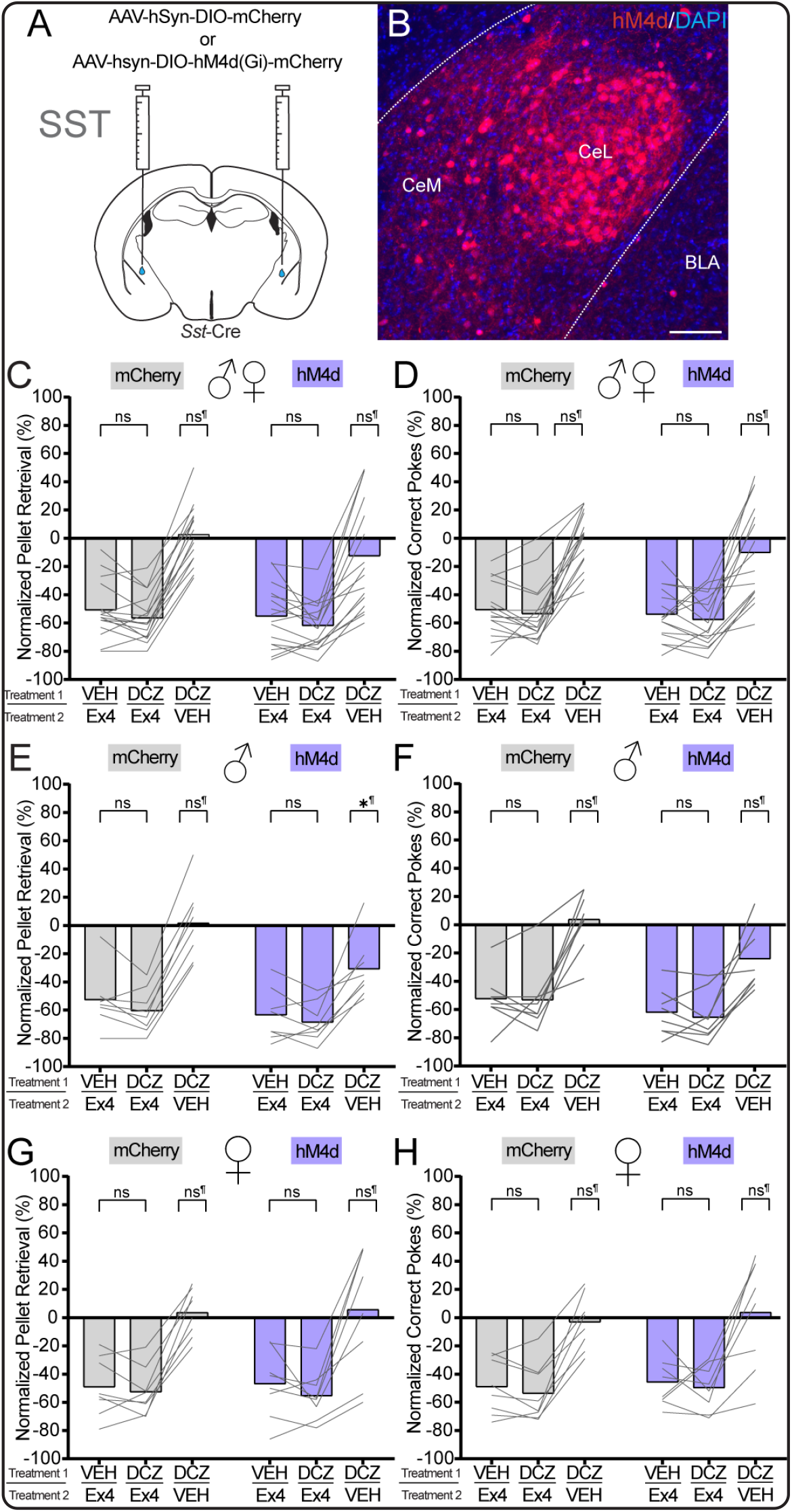
Chemogenetic inhibition of *Sst*^CeA^ neurons has no effect on systemic Exendin-4-mediated hypophagia. **A.**Surgical schematic of bilateral injection of either AAV-hSyn-DIO-mCherry (n = 8/sex) or AAV-hSyn-hM4d-mCherry (n = 8/sex) in CeA for *Sst*-Cre mice. **B**. Representative image a hM4d-mCherry-expressing tissue from Sst-Cre mouse taken at 20x magnification. Scale bar = 100 µM. **C**. Pellets retrieved by all *Sst*-Cre mice during each treatment (Tx) session. Gr X Tx: F(1.893, 56.79) = 1.333, *p* = 0.2712; Gr effect: F(1, 30) = 1.295, *p* = 0.2641; Tx effect: F(1.893, 56.79) = 124.6, *p* < 0.0001. Post tests: Veh/Ex4 vs DCZ/Ex4: mCh (*p* = 0.2854), hM4 (*p* = 0.3097); DCZ/Veh vs Veh/Veh: mCh (*p* =0.9548), hM4 (*p* = 0.5872). **D**. Active port pokes by all *Sst*-Cre mice during each Tx session. Gr X Tx: F(2.322, 69.66) = 0.6302, *p* = 0.5585; Gr effect: F(1, 30) = 0.8424, *p* = 0.3660; Tx effect: F(2.322, 69.66) = 114.0, *p* < 0.0001. Post tests: Veh/Ex4 vs DCZ/Ex4: mCh (*p* = 0.8651), hM4 (*p* = 0.8351); DCZ/Veh vs Veh/Veh: mCh (*p* = 0.9998), hM4 (*p* =0.6391). **E**. Pellets retrieved by male *Sst*-Cre mice during each Tx session. Gr X Tx: F(2.214, 31.00) = 4.811, *p* = 0.0128; Gr effect: F(1, 14) = 3.963, *p* = 0.0664; Tx effect: F(2.214, 31.00) = 100.3, *p* < 0.0001. Post tests: Veh/Ex4 vs DCZ/Ex4: mCh (*p* = 0.3220), hM4 (*p* = 0.6511); DCZ/Veh vs Veh/Veh: mCh (*p* = 0.9972), hM4 (*p* = 0.0199). **F**. Active port pokes by male *Sst*-Cre mice during each Tx session. Gr X Tx: F(3, 42) = 2.583, *p* = 0.0793; Gr effect: F(1, 14) = 4.338, *p* = 0.0561; Tx effect: F(2.469, 34.56) = 73.58, *p* < 0.0001. Post tests: Veh/Ex4 vs DCZ/Ex4: mCh (*p* = 0.9993), hM4 (*p* = 0.8733); DCZ/Veh vs Veh/ Veh: mCh (*p* = 0.9531), hM4 (*p* = 0.0686). **G**. Pellets retrieved by female *Sst*-Cre mice during each Tx session. Gr X Tx: F(1.899, 26.59) = 0.07238, *p* = 0.9277; Gr effect: F(1, 14) = 0.002488, *p* = 0.9609; Tx effect: F(1.899, 26.59) = 50.50, *p* < 0.0001. Post tests: Veh/Ex4 vs DCZ/Ex4: mCh (*p* = 0.8671), hM4 (*p* = 0.5803); DCZ/Veh vs Veh/Veh: mCh (*p* = 0.9259), hM4 (*p* = 0.9829). **H**. Active port pokes by female *Sst*-Cre mice during each Tx session. Gr X Tx: F(2.112, 29.56) = 0.1134, *p* = 0.9026; Gr effect: F(1, 14) = 0.2487, *p* = 0.6257; Tx effect: F(2.112, 29.56) = 46.86, *p* < 0.0001. Post tests: Veh/Ex4 vs DCZ/ Ex4: mCh (*p* = 0.5993), hM4 (*p* = 0.9575); DCZ/Veh vs Veh/Veh: mCh (*p* = 0.9669), hM4 (*p* = 0.9926). For C-H, data were analyzed using two-way repeated measures ANOVA with Tukey’s multiple comparisons. Datapoints for each mouse during each Tx session were normalized to their respective Veh/Veh session. ¶ Indicates a post test to Veh/Veh condition. Raw data in Figure S3.

Ultimately, these data demonstrate that activation of *Sst*^CeA^ neurons is not required for the complete hypophagic effect of Ex-4.

### 2.5 Inhibition of *Glp1r*^CeA^ neurons rescues GLP-1R agonist-induced hypophagia

Previously, our lab has shown that GLP-1Rs are expressed in the CeA, enriched within the medial division of the CeA (CeM), and do not overlap with other genetic markers like Somatostatin and PKCδ [54]. Exogenous administration of GLP-1R agonists consistently activates the CeA and decreases food intake, but whether *Glp1r*^CeA^ neurons are necessary to mediate GLP-1R agonist-induced hypophagia has not been tested [75]. We bilaterally injected mCherry or hM4d into the CeA of *Glp1r*-Cre mice (**Figure 5A**), resulting in robust mCherry expression within the CeM as we previously demonstrated (**Figure 5B**). In control animals, DCZ treatment prior to Ex-4 resulted in no change in pellets retrieval and active pokes relative to Veh/Ex-4 conditions (**Figure 5C-D, S4A-B**). In hM4d animals; however, we observed a significant increase of pellet retrieval and a non-significant increase of active pokes relative to Veh/Ex-4 conditions in their normalized data (**Figure 5C-D**). Similarly, raw pellet retrieval and active pokes were non-significantly increased (**S4A-B**). When grouped data were parsed by sex, we observed a nonsignificant increase in male (**Figure 5E-F, S4C-D**) and female (**Figure 5G-H, S4E-F**) hM4d mice. When examining the effects of DCZ/Veh alone, *Glp1r*^CeA^ neuron inhibition did not significantly alter pellet retrieval and active pokes of hM4d mice (**Figure 5C-H, S4A-F**) relative to Veh/Veh. Collectively, these data demonstrate that activation of *Glp1r*^CeA^ neurons is required for the complete hypophagic effect of Ex-4.

**Figure 5.**
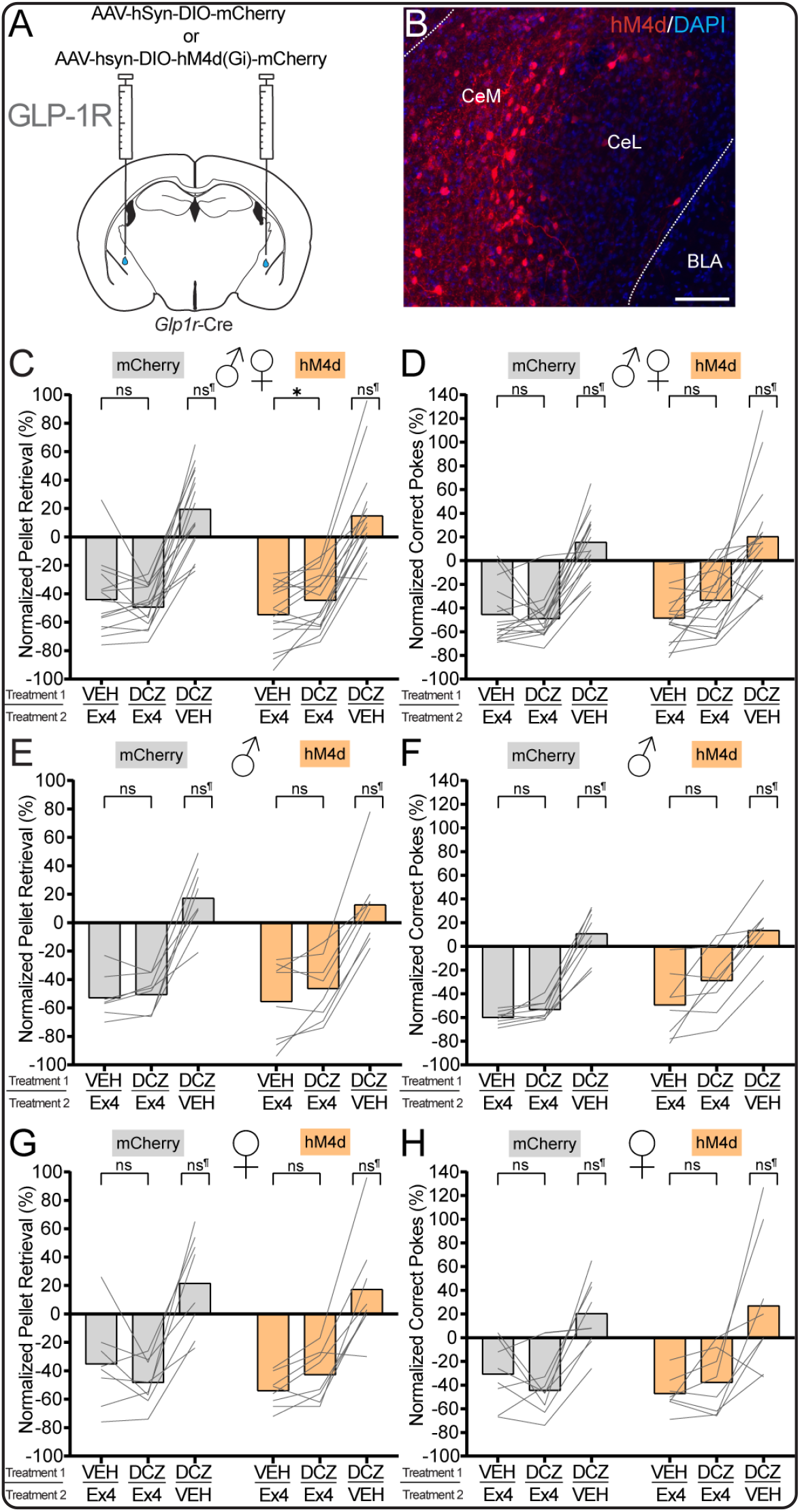
Chemogenetic inhibition of *Glp1r*^CeA^ neurons attenuates systemic Exendin-4-mediated hypophagia. **A.**Surgical schematic of bilateral injection of either AAV-hSyn-DIO-mCherry (n = 8/sex) or AAV-hSyn-hM4dmCherry (n = 8/sex) the in CeA of *Glp1r*-Cre mice. **B**. Representative image of hM4d-mCherry expressing CeA tissue from *Glp1r*-Cre mouse taken at 20x magnification. Scale Bar = 200 µM. **C**. Pellets retrieved by all *Glp1r*-Cre mice during each treatment (Tx) session. Group (Gr, mCherry/hM4d) X Tx: F(2.194, 65.83) = 1.148, *p* = 0.3270; Gr effect: F(1, 30) = 0.2528, *p* = 0.6188; Tx effect: F(2.194, 65.83) = 118.2, *p* < 0.0001. Post tests: Veh/Ex4 vs DCZ/ Ex4: mCh (*p* = 0.6219), hM4(*p* = 0.0290); DCZ/Veh vs Veh/Veh: mCh(*p* = 0.0644), hM4 (*p* = 0.2987). **D**. Active port pokes by all *Glp1r*-Cre mice selected during each Tx session. Gr X Tx: F(2.155, 64.66) = 1.107, *p* = 0.3400; Gr effect: F(1, 30) = 0.6597, *p* = 0.4231; Tx effect: F(2.155, 64.66) = 67.79, *p* < 0.0001. Post tests: Veh/Ex4 vs DCZ/Ex4: mCh(*p* = 0.9937), hM4(*p* = 0.1459); DCZ/Veh vs Veh/Veh: mCh(*p* = 0.1627), hM4(*p* = 0.4084). **E**. Pellets retrieved by male *Glp1r*-Cre mice during each Tx session. Gr X Tx: F(2.311, 32.35) = 0.2455, *p* = 0.8140; Group effect: F(1, 14) = 0.01258, *p* = 0.9123; Tx effect: F(2.311, 32.35) = 80.73, *p* < 0.0001. Post tests: Veh/Ex4 vs DCZ/Ex4: mCh(*p* = 0.8308), hM4(*p* = 0.4447); DCZ/Veh vs Veh/Veh: mCh(*p* = 0.2264), hM4(*p* = 0.6472). **F**. Active port pokes by male *Glp1r*-Cre mice during each Tx session. Gr X Tx: F(2.689, 37.65) = 2.263, *p* = 0.1030; Gr effect: F(1, 14) = 2.070, *p* = 0.1722; Tx effect: F(2.689, 37.65) = 79.00, *p* < 0.0001. Post tests: Veh/Ex4 vs DCZ/Ex4: mCh(*p* = 0.1063), hM4(*p* = 0.2277); DCZ/Veh vs Veh/Veh: mCh(*p* = 0.5166), hM4(*p* = 0.4588). **G**. Pellets retrieved by female *Glp1r*-Cre mice during each Tx session. Gr X Tx: F(1.972, 27.61) = 1.095, *p* = 0.3479; Gr effect: F(1, 14) = 0.3242, *p* = 0.5781; Tx effect: F(1.972, 27.61) = 42.94, *p* < 0.0001. Post tests: Veh/Ex4 vs DCZ/Ex4: mCh(*p* = 0.4631), hM4(*p* = 0.1093); DCZ/Veh vs Veh/Veh: mCh(*p* = 0.3607), hM4(*p* = 0.5846). **H**. Active port pokes by female *Glp1r*-Cre mice during each Tx session. Gr X Tx: F(1.747, 24.45) = 0.6367, *p* = 0.5171; Gr effect: F(1, 14) = 0.009802, *p* = 0.9225; Tx effect: F(1.747, 24.45) = 21.74, *p* < 0.0001. Post tests: Veh/Ex4 vs DCZ/Ex4: mCh(*p* = 0.6059), hM4(*p* = 0.6198); DCZ/Veh vs Veh/Veh: mCh(*p* = 0.3114), hM4(*p* = 0.5861). For C-H, data were analyzed using two-way repeated measures ANOVA with Tukey’s multiple comparisons. Datapoints for each mouse during each Tx session were normalized to their respective Veh/Veh session. ¶ Indicates a post test to Veh/Veh condition. Raw data in Figure S4.

### 2.6 Inhibition of *Glp1r*^CeA^ neurons attenuates the effect of Ex-4 on palatable and energy-dense food

In a previous study, we demonstrated a population of prepronociceptin-expressing neurons in the CeA was important for the consumption of energy-dense and highly palatable food [66]. Further, one of the hallmark effects of GLP-1R agonists is their ability to reduce preference for and consumption of energy-dense, palatable foods [12], [81], [82]. Thus, we reasoned that *Glp1r*^CeA^ neurons might be tuned to mediate suppression of palatable high-fat diet (HFD) by Ex-4. To test this hypothesis, we generated new mice using the same approach used in Figure 5. As FED3 is incompatible with the use of HFD, we used standard HFD pellets weighed manually. To avoid entrainment, *ad libitum* chow fed mice were allowed access to HFD once every four days in combination with chow but receiving randomized two drug treatments as described in Figure 2. In this way, we used a model of hedonic hyperphagia to test the contribution of *Glp1r*^CeA^ neurons to the effects of exogenous Ex-4 (**Figure 6A-B**). In mCherry control animals, 5 µg/kg Ex-4 produced a ∼60% reduction in HFD intake while also increasing the relative percent of chow during this access period (**Figure 6C-D, S5A**). We observed no effect of DCZ pre-treatment in this group on Ex-4 or on its own. In contrast, DCZ pretreatment of the hM4d group produced a highly significant attenuation of the hypophagic effects of Ex-4 on HFD intake (**Figure 6C, S5A**). Although a decreasing trend was observed, DCZ did not significantly rescue the increase in percent chow intake or decreased HFD preference seen with Ex-4. When separated by sex, we observed similar phenomena. Chemogenetic inhibition of *Glp1r*^CeA^ neurons produced a significant attenuation of Ex-4’s suppression of HFD intake without producing a significant rescue of HFD preference in both male and female mice (**Figure 6E-H, S5B-C**). These data demonstrate that activation of *Glp1r*^CeA^ neurons is required for the complete suppression of palatable food intake produced by Ex-4

**Figure 6.**
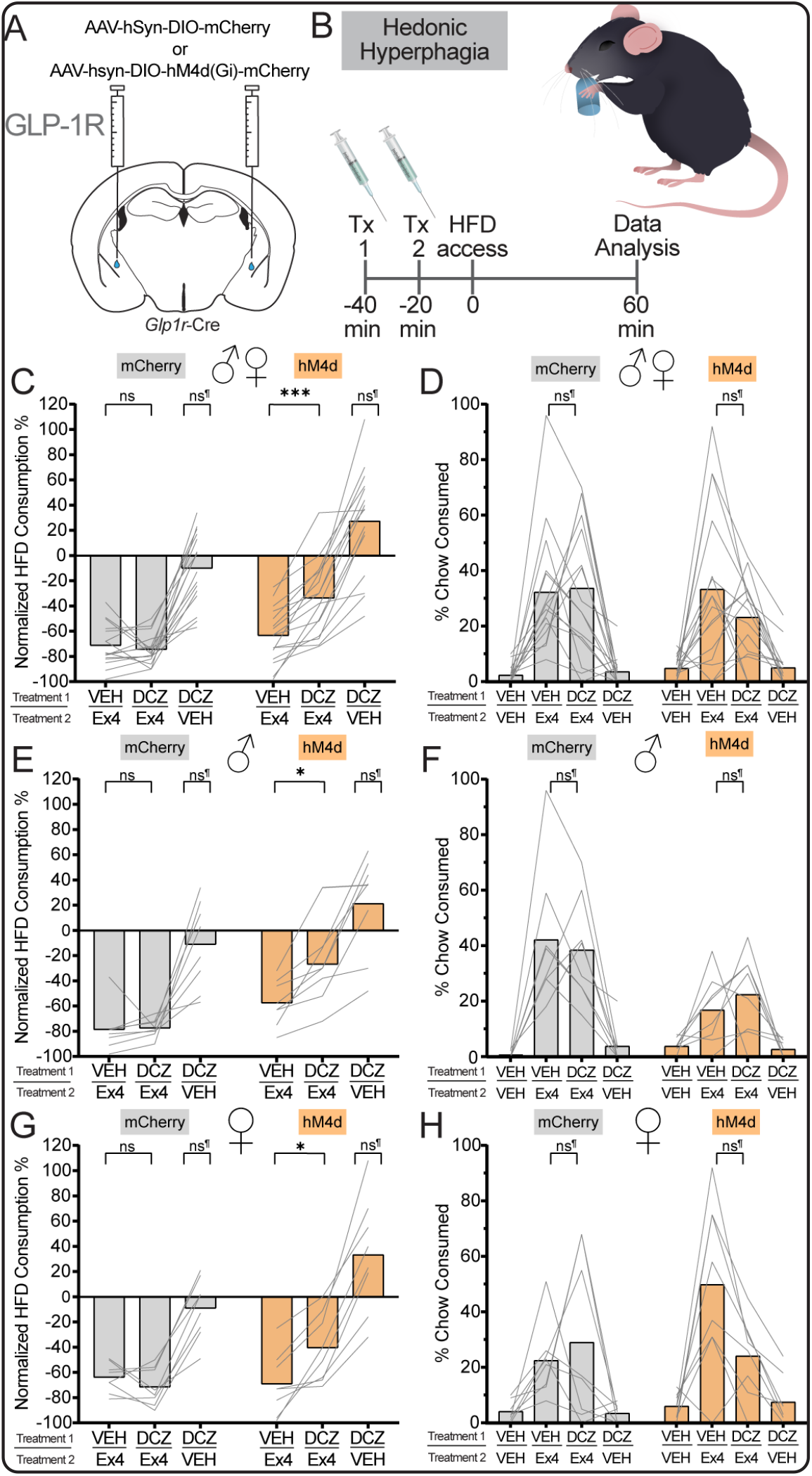
Chemogenetic inhibition of *Glp1r*^CeA^ neurons attenuates systemic Exendin-4-mediated reduction in palatable food intake. **A.** Surgical schematic of bilateral injection of either AAV-hSyn-DIO-mCherry (n = 8/sex) or AAV-hSyn-hM4d-mCherry (n = 8/sex) into the CeA of Glp1r-Cre mice. B. Representative illustration of timeline for each intermittent high-fat diet (HFD) access/treatment (Tx) session (i.p.) and endpoint (60 min) for data acquisition for analysis. Each animal is exposed to each Tx regime (Fig. 2F) at least once over the course of 4 sessions. C. Relative percentage of HFD consumed by all Glp1r-Cre mice during each Tx session. Group (mCherry/hM4d) X Tx: F(1.969, 59.07) = 8.847, p = 0.0005; Group effect: F(1, 30) = 14.86, p = 0.0006; Tx effect: F(1.969, 59.07) = 120.9, p < 0.0001. Post tests: Veh/Ex4 vs DCZ/Ex4: mCherry (p = 0. 0.9742), hM4d (p = 0.0001); DCZ/Veh vs Veh/Veh: mCherry (p = 0.6910), hM4d (p = 0.1105). D. Percent of chow consumed by all Glp1r-Cre mice selected during each Tx session. Group X Tx: F(1.972, 59.15) = 1.221, p = 0.3019; Group effect: F(1, 30) = 0.1699, p = 0.6831; Tx effect: F(1.972, 59.15) = 31.78, p < 0.0001. Post tests: Veh/Ex4 vs DCZ/Ex4: mCherry (p = 0.9977), hM4d (p = 0.4654); DCZ/Veh vs Veh/Veh: mCherry (p = 0.8345), hM4d (p = 0.9997). E. Relative percentage of HFD consumed by male Glp1r-Cre mice during each Tx session. Group X Tx: F(1.922, 26.91) = 4.252, p = 0.0261; Group effect: F(1, 14) = 11.15, p = 0.0049; Tx effect: F(1.922, 26.91) = 51.52, p < 0.0001. Post tests: Veh/Ex4 vs DCZ/Ex4: mCherry (p = 0.9999), hM4d (p = 0.0288); DCZ/Veh vs Veh/Veh: mCherry (p = 0.9490), hM4d (p = 0.6828). F. Percent of chow consumed by male Glp1r-Cre mice during each Tx session. Group X Tx: F(2.072, 29.00) = 4.412, p = 0.0202; Group effect: F(1, 14) = 6.603, p = 0.0223; Tx effect: F(2.072, 29.00) = 25.02, p < 0.0001. Post tests: Veh/Ex4 vs DCZ/Ex4: mCherry (p = 0.9657), hM4d (p = 0.8740); DCZ/Veh vs Veh/Veh: mCherry (p = 0.6030), hM4d (p = 0.8750). G. Relative percentage of HFD consumed by female Glp1r-Cre mice during each Tx session. Group X Tx: F(1.739, 24.34) = 6.232, p = 0.0085; Group effect: F(1, 14) = 4.157, p = 0.0608; Tx effect: F(1.739, 24.34) = 71.24, p < 0.0001. Post tests: Veh/Ex4 vs DCZ/Ex4: mCherry (p = 0.7839), hM4d (p = 0.0235); DCZ/Veh vs Veh/Veh: mCherry (p = 0.8976), hM4d (p = 0.3790). H. Percent of chow consumed by female Glp1r -Cre mice during each Tx session. Group X Tx: F(1.994, 27.92) = 2.987, p = 0.0669; Group effect: F(1, 14) = 2.269, p = 0.1542; Tx effect: F(1.994, 27.92) = 14.70, p < 0.0001. Post tests: Veh/Ex4 vs DCZ/Ex4: mCherry (p = 0.9557), hM4d (p = 0.0638); DCZ/Veh vs Veh/Veh: mCherry (p = 0.9774), hM4d (p = 0.9803). For C-H, data were analyzed using two-way repeated measures ANOVA with Tukey’s multiple comparisons. For C, E, G, datapoints for each mouse during each Tx session were normalized to their respective Veh/ Veh session. For D, F, H, percentage of chow consumed was calculated by dividing the consumed gram amounts of standard chow pellet by the summation of standard chow pellet and HFD pellet in respective to each animal for each Tx session. ¶ Indicates a post test to Veh/Veh condition. Raw data in Figure S5.

## 3. Discussion

In this study, we observed that peripheral administration of Ex-4 increases CeA neuronal activity in freely behaving mice and that this was blocked by a systemic GLP-1R antagonist. Importantly, we observed heterogenous responses across animals and this may be due to fiber placement and/ or recording of specific subpopulations. Further efforts are needed to reveal the specific cell types activated by Ex-4 or other GLP-1R compounds and the mechanisms, direct or indirect, by which this activation occurs through GLP-1R.

At a behavioral level, chemogenetic inhibition of total and select CeA neuronal populations increased feeding in the presence of systemic Ex-4, however; GLP-1R agonist induced hypophagia was incompletely rescued. It is likely that the engagement of parallel anorexigenic circuitry throughout the CNS continues to mediate GLP-1R signaling despite silencing the CeA, as well as the potential contribution of uncharacterized CeA cell types. Indeed, the partial attenuation observed during complete CeA inhibition underscores that the CeA operates as a critical node within a broader, brain-wide anorexigenic system. Gabery and others have shown that systemic GLP-1R agonists directly activate the hindbrain as shown by the induction of Fos and accumulation of fluorescently-conjugated semaglutide. They also observed activation of the CeA and other brain regions but were unable to detect fluorescently-conjugated semaglutide. These data suggest that there exist primary and secondary sites of activation in response to peripheral GLP-1R agonists, which is consistent with the anatomical wiring of the CeA positioning it to integrate and translate upstream GLP-1R signals into coordinated behavioral outputs, supported by our data [12], [22], [75]. In contrast, Qiao and others reported that an electrical lesion of the CeA in male rats minimally attenuated the hypophagic effects of Ex-4 on homeostatic chow intake but did attenuate high sucrose consumption following a 24hr fast [57]. We speculate that a CeA lesion likely disrupts fibers of passage and eliminates multiple opposing populations that promote and constrain the actions of peripheral GLP-1R agonists. In support of this notion, it is known that *Prkcd*^CeA^ and *Sst*^CeA^ neurons inhibit one another to regulate fear learning [80].

Beyond parallel circuitry, the population specificity of our findings suggests that the CeA itself integrates exogenous GLP-1R signaling through heterogeneous and functionally distinct neuronal populations. The robust attenuation of Ex-4 hypophagia by *Prkcd*^CeA^ inhibition indicates that up-stream projections likely converge on the CeA along with parallel GLP-1R circuitry in other CNS nuclei. This is consistent with *Prckd*^CeA^ neurons being known to mediate anorexigenic signals in general [78]. Additionally, that *Sst*^CeA^ inhibition produced no significant attenuation of Ex-4 induced hypophagia highlights the specificity of these effects. The opposing relationship between *Prkcd*^CeA^ and *Sst*^CeA^ neurons as demonstrated for fear learning and pain encoding raises the possibility that intact reciprocal inhibition within the CeA is important for how GLP-1R signals are processed and translated into behavioral outputs. Silencing one population may disinhibit the other, potentially explaining why full rescue was never achieved. Taken together, these findings suggest that exogenous GLP-1R signaling engages the CeA through select neuronal populations, particularly *Prkcd*^CeA^ neurons, and likely coordinates with downstream nuclei to mediate the complete hypophagic response to peripheral GLP-1R agonist administration.

In these studies, we examined differences in food consumption during chemogenetic inhibition of the CeA neuronal populations fully powered to detect sex differences. Depending on the population of interest, we either detected or failed to detect sex differences. Across species, there are sex differences towards feeding attributable to endogenous hormones such as estrogens and androgens [83]. Further, several studies have demonstrated interactions between GLP-1R and estradiol signaling, where exogenous GLP-1R agonists produce stronger hypophagic effects in females in a estrus cycle-dependent manner [38], [84], [85]. In our studies, we did not account for the estrus cycle in female subjects and this may have impacted responses to Ex-4 during chemogenetic inhibition and the observed variability in some cases, particularly when inhibiting whole CeA. However, there is a notable contrast with selectively inhibition of *Prkcd*^CeA^ neurons in female mice, where the attenuation of Ex-4 induced hypophagia is robust relative to both *vGat*^CeA^ male and female mice and even *Prkcd*^CeA^ male mice. We speculate that select neuronal populations may be less sensitive to cycle-dependent hormonal signaling. Further, studies of this design would benefit from carefully tracking estrus cycle or including ovariectomized females in combination with estrogen analog treatments.

Chemogenetic inhibition of *Glp1r*^CeA^ neurons produced a notably stronger attenuation of Ex-4 hypophagia on HFD consumption(30%) compared to standard grain-based diet(10%). This suggests that *Glp1r*^CeA^ neurons may play a more prominent role in the consumption of palatable, energy-dense diets than in homeostatic feeding of calorically balanced diets. Importantly, when animals were presented with both HFD and standard chow during intermittent HFD assays, Ex-4 decreases HFD preference resulting in more chow intake but this phenomenon trended lower in both sexes combined and in females specifically. This indicates that the rescue of feeding observed during *Glp1r*^CeA^ inhibition was not just an increase in appetite but also shifted preference back towards palatable, energy-dense diet consumption. Consistent with a recent preprint, this behavioral evidence suggests that *Glp1r*^CeA^ neurons may suppress hedonic-driven rather than homeostatic feeding [86]. Overall, this distinction may be due to differences in diet palatability. In a previous study, we demonstrated that *Pnoc*^CeA^ neurons are activated following intermittent HFD access and specifically promote palatable food consumption [66]. Given that inhibiting *Glp1r*^CeA^ neurons attenuates Ex-4 induced hypophagia while *Pnoc*^CeA^ neurons can promote palatable, energy-dense food consumption, we speculate that these two populations may function in a reciprocal manner within the CeA to bidirectionally regulate hedonic feeding. This potential relationship warrants investigation in future studies and to determine the mechanisms behind differences in diet effects.

Our findings capture the acute contribution of *Glp1r-* ^CeA^ neurons towards Ex-4 induced hypophagia; however, the chronic physiological role of these neurons in feeding and energy homeostasis has yet to be determined. Across other CNS systems, chronic silencing or ablation of GLP-1R expressing neuronal reveals homeostatic contributions that acute manipulation alone does not fully capture. For instance, in the arcuate nucleus (ARC), genetic ablation of GLP-1R expressing neurons fails to attenuate the hypophagic effect of systemically administered semaglutide; however, chronic silencing of thyrotropin-releasing hormone-expressing neurons that co-express GLP-1Rs within the ARC does partially attenuate the hypophagic and weight loss effects of acutely and chronically systemic administration of liraglutide [21], [87]. Within the dorsomedial hypothalamus, viral knockdown of GLP-1R expressing neurons fails to produce hyperphagia but did decrease energy expenditure and attenuated Ex-4 induced hypophagia [36]. Of the limbic sites, the lateral septum has abundant GLP-1R expression, where knockdown of GLP-1R expressing neurons within the LS attenuates the hypophagic and weight loss effects of liraglutide over time and chronically silencing these neurons acutely attenuates liraglutide’s hypophagic effects, suggesting that limbic GLP-1R populations may have modulatory roles more clearly revealed under chronic paradigms [15]. Whether chronic silencing or ablation of *Glp1r*^CeA^ neurons can produce similar outcomes warrants further investigation given that clinical usage of GLP-1R agonists are administered chronically. Further studies that incorporate targeted silencing or ablation of *Glp1r*^CeA^ neurons will be important to elucidate their role in energy balance and contributions to chronic GLP-1R agonism.

Although it is clear that 1) the majority of peripherally administered GLP-1R agonist will accumulate in circumventricular organs of the brain and 2) above we discuss the evidence that suggests a pathway for indirect activation of the CeA by systemic GLP-1R agonists, our data with *Glp1r*^CeA^ neurons suggest that it is <u>possible</u> that Ex-4 or other peripherally-administered GLP-1R agonists may directly activate *Glp1r*^CeA^ neurons. The posterior CeA has close proximity to the lateral ventricles and previous whole-brain imaging approaches using fluorescently-conjugated liraglutide and semaglutide may not resolve low-level binding. Consistent with this idea, Goddschall *et al* have demonstrated that peripheral administration of the small molecule GLP-1R agonists, danuglipron and orforglipron, can activate the CeA expressing humanized GLP-1Rs [86]. Whether GLP-1R expression levels in the CeA are sufficient for direct engagement by GLP-1R agonists such as Ex-4 and various other exogenous agonists has yet to be fully characterized. Collectively, we speculate that CeA activation by peripheral GLP-1R agonists is likely multifaceted, potentially incorporating both indirect and direct signaling. Future studies utilizing site specific methods and rigorous treatment regimens with doses of multiple GLP-1R compounds will be critical to help elucidate this model.

We aimed to rigorously perform and analyze the data present in this study; however, we acknowledge several limitations. First, while DCZ is generally regarded pharmacologically inert in animals lacking transduction of DREADDs in targets of interest, it is not without off-target effects [77]. We selected DCZ over clozapine-N-oxide (CNO) due to its reported minimal off-target effects and ultimately reported no significant changes with any of the mCherry mice used for FR1 refeed and intermittent HFD access except for female *vGat*^CeA^ mice during assessment of active pokes, where we report a significant decrease. The basis for this finding is unclear. This finding complicates the interpretation of the female *vGat*^CeA^ specifically, as these findings were not replicated in other mCherry-expressing transgenic strains. Given that DCZ is relatively recent compared to CNO which is still widely used, future chemogenetic studies should take into account for potential off-target effects of DCZ. Second, while viral transduction was predominantly contained within the CeA, we can not rule occasional spread to adjacent regions. Third, the present study did not directly assess the role of GLP-1R protein signaling. While our findings demonstrate that select populations are necessary for the full hypophagic response of peripherally administered Ex-4, whether GLP-1R is directly engaged in these neurons or whether recruitment occurs indirectly in upstream circuitry remains to be determined.

For the first time, we show that chemogenetic inhibition of CeA neurons attenuates the complete hypophagic effects of Ex-4 on food intake and food-seeking. Additionally, in a similar manner, population-specific inhibition revealed varying contributions to Ex-4 mediated hypophagia. Our data demonstrate that inhibition of all CeA neurons, *Prkcd*^CeA^ neurons, or *Glp1r*^CeA^ neurons produced a significant attenuation of Ex-4 mediated hypophagia. Inhibition of *Sst*^CeA^ neurons, on the other hand, had no effect on Ex-4 mediated hypophagia. These data demonstrate multiple, non-overlapping neuron populations in the CeA participate in mediating the actions of exogenous GLP-1R agonists.

## 4.1 Materials and Methods

### 4.1 Animals

We used 8-16 week old C57BL6/J, *Slc32a1*^*tm2*(cre)Lowl^ (vGat; MGI: 5751790), Tg(*Prkcd*-glc-1/CFP,-cre)EH124Gsat (PC; MGI: 3844446), *Sst*^*tm2*.*1-Cre*^ (SC; MGI: 4838416) and *Glp1r*^tm1.1(cre) Lbrl^ (GL; MGI: 5776617) male and female mice throughout this study (Jackson Labs, Bar Harbor, ME) [88], [89], [90], [91]. Unless otherwise specified, mice were group housed with 3-5 mice per cage on standard bedding and provided with food and water *ad libitum*. A 12:12 light cycle was maintained with lights on at 7AM and off at 7PM. All procedures were performed with approval from the University of Alabama at Birmingham Institutional Care and Use Committee (IACUC).

### 4.2 Histology and Fluorescence Immunohistochemistry

A lethal dose of 2,2,2-tribromoethanol (250 mg/kg, Sigma-Aldrich) was administered via intraperitoneal injection prior to sacrifice. After an appropriate anesthetic depth was confirmed by loss of the toe pinch reflex, animals were transcardially perfused with 50 ml of 0.01 M PBS, then 50 ml of 4% paraformaldehyde (Electron Microscopy Sciences, Cat No. 19210) in 0.01 M PBS. Brains were extracted, post-fixed in 4% paraformaldehyde overnight then transferred to 30% sucrose (Thermo Fischer Scientific, Cat No. 036508.A1) for 36-48 hr. Once sufficiently cryoprotected, brains were sectioned at 40 µm on a Leica CM1850 cryostat. Free-floating coronal tissue sections were washed three times in 0.01 M PBS for 5 min each, then permeabilized for 30 min in 0.5% Triton X-100/PBS (Fisher Scientific, Cat No. BP151-500), followed by another three washes. Sections were then incubated in a blocking solution comprised of 0.1% Triton X-100/PBS, 1% Bovine Serum Albumin (Sigma, Cat No. A2153), and 10% Normal Donkey serum (Jackson ImmunoResearch, Cat No. 017-000-121) for 1 hr. Following blocking, sections were then transferred into a separate blocking solution containing rabbit anti-mCherry polyclonal antibody (1:500; Invitrogen, Thermo Fisher Scientific Cat No. PA5-34974) and incubated overnight at 4 °C. The following day, sections were then washed three times in 0.01 M PBS for 5 min each. Sections were then incubated for 2 h in PBS containing donkey anti-rabbit conjugated with Cy™3 AffiniPure® (1:200; Jackson ImmunoResearch, Cat No. 711-165-152), then washed three times in 0.01 M PBS for 5 min each again. Once completed, sections were then mounted on SuperFrost slides (Fisher Scientific, Cat No. 12-550-123), allowed to dry, then coverslipped with Vectashield Hard-set mounting medium with DAPI (Vector Labs, Cat No. H-1500-10).

### 4.3 Stereotaxic Surgery

Surgical procedures were performed on 6–8-week-old male and female mice. Animals received Buprenorphine SR (1.0 mg/ kg, s.c.) and Meloxicam SR (5 mg/kg, s.c.) prior to surgery and again on post-op day 2. Mice were anesthetized with isoflurane inhalation (0.5 −5%) and secured in a stereotaxic frame. Eye lubricant was applied immediately and as needed throughout surgery. The respiratory rate was visually monitored throughout surgery with a target of 55-60 breaths/minute. Internal temperature was maintained at 35°C using a homoeothermic biodynamic feedback system (Harvard Apparatus; Holliston, MA). Anesthetic depth was confirmed by a loss of toe pinch reflex. The scalp was depilated and sterilized with rotating application of 70% ethanol and betadine. Triple antibiotic ointment and 4% topical lidocaine ointment was also applied. A midline incision was made and the scalp parted to expose the skull surface. After verifying appropriate orientation of the head, 1 mm burr holes were drilled for injection/fiber placement and screw stabilization. For viral injections, we used a pulled glass capillary secured in Nanoject III (Drummond Scientific; Broomall, PA). Viruses for injection were either AAV-hSyn-DIO-mCherry (Addgene, Cat No. 50459-AAV9), AAV-hSyn-DIO-hM4d(Gi)-mCherry (Addgene, Cat No. 44362-AAV9), pGP-AAV-syn-FLEX-jGCaMP7f-WPRE (Addgene, Cat No. 104492-AAV9) and were tittered from stock concentration into sterile PBS at the following concentrations respectively, 6.25*1012 GC/ml, 5.75*10^12^ GC/ml, and 2.8*10^13^ GC/ml. For CeA, 100-300 nl of each virus was injected at +/-3.0mm lateral, −1.3mm posterior, −4.6 mm ventral of bregma. For virus-only injected animals, we closed the scalp using veterinary adhesive (3M, Cat No. 1469SB). For mice used for photometry, optical fibers (Newdoon, Cat No. FOCC-B-200-1.25-0.37-10) were secured in a custom-made ferrule holder (https://github.com/OpenBehavior/Modular-Stereotaxic-Holders) and implanted at a 5° angle +/-3.05 mm lateral, - 1.3 mm posterior, −4.45 mm ventral of bregma. Fibers were secured with setscrews and Metabond (Parkell, Cat No. S380) applied over the skull surface, covering the base of the fibers and the screws. When the Metabond was dry, dental cement (Stoelting Co., Cat No. 10-000-786) was applied and allowed to harden before rehousing the animal. Virally injected animals were re-housed with their littermates and fiber implanted animals were singly housed. Post-operative care included daily weight monitoring for 5 days or until their weight returned to within 5% of their initial preoperative weights. All mice recovered for 3 weeks under normal husbandry conditions to allow for viral expression and recovery prior to any experiment.

### 4.4 Subject Inclusion Criteria

Validation of cells transduced by Cre-dependent viruses in the CeA and the termination sites of fiber optic implants above the CeA were visually confirmed first under an epifluorescent microscope and then a Keyence BZ-X810 Fluorescent Microscope and then noted within written records. For photometry experiments, CeA hemispheres that either did not have transduced cells in the CeA or did not have their fiber optic directly above the CeA were excluded from statical analyses. For feeding-related experiments, animals that did not have expression of transduced cells present within the CeA and in both hemispheres were excluded from our statical analyses. For animals feeding from FED3s, they needed to sufficiently learn to feed from their FED3 under an FR1 paradigm; meaning that following a fasted condition and prior to any treatment regimes, they needed to successfully poke the active port, retrieve a pellet, and consume at least 0.2 grams (10 precision pellets) within the 1st hour to ensure they are trained for refeeding experiments. Mice that failed to learn to feed from their FED3 or displayed pellet-hoarding behaviors anytime throughout FR1 feeding assays were excluded from our studies. Pellet hoarding is defined as pellets being successfully being retrieved from the FED3 but were dropped and not consumed during experimental treatments. For intermittent HFD access feeding studies, mice also needed to consume atleast 0.2 grams of the HFD pellet within the 1st hour following saline injections prior to any treatment regimes. Mice that failed to consistently consume this amount were excluded from statical analyses.

### 4.5 Electrophysiology

#### 4.5.1 Slice preparation

Electrophysiological experiments were performed as previously described [54]. Briefly, animals were removed from their cage and brought to the lab for brain slice preparation. The animal rested in a quiet chamber for 30 min prior to slice preparation to dissipate stress associated with animal transport from the vivarium. Mice were treated with a lethal dose of 2,2,2-tribromoethanol (250 mg/kg, i.p.), and, after a deep plane of anesthesia was reached, animals were transcardially perfused with cold, sodium free N-methyl-D-glucamine (NMDG) artificial cerebrospinal fluid (aCSF)[(in mM) 93 N-methyl-D-glucamine, 2.5 KCl, 1.2 NaH2PO4, 30 NaHCO3, 20 HEPES, 25 Glucose, 5 L-ascorbic acid, 2 Thiourea, 3 sodium pyruvate, 10 MgSO4 X 7H2O, 0.5 CaCl2 X 2H2O]. All solutions were saturated with 95% O2 and 5% CO2. The brain was rapidly dissected and coronal 300 µM sections prepared in ice cold, oxygenated NMDG aCSF using a Leica VT1200S at 0.07 mm/s. Slices were immediately transferred to 36°C NMDG aCSF for 10 min, and then normal 36°C aCSF [(in mM): 124 NaCl, 4.4 KCl, 2 CaCl2, 1.2 MgSO4, 1 NaH2PO4, 10.0 glucose, and 26.0 NaHCO3]. Slices rested in normal aCSF for at least 30 min prior to recordings.

#### 4.5.2 Recordings

Whole cell or cell-attached patch clamp recordings were performed in the CeA guided by DIC microscopy and mCherry fluorescence. Slices were then transferred to a recording chamber (Warner Instruments), submerged in normal, oxygenated aCSF and maintained at 32°C with a flow rate of 2 ml/min. For determining hM4d expression, we patched mCherry-expressing cells in the CeA. For hM4d recordings we used a potassium gluconate internal solution [(in mM): 135 C6H11KO7, 5 NaCl, 2 MgCl2, 20 HEPES, 0.6 EGTA, 4 Na2ATP, 0.4 Na2GTP at a final osmolarity of 290 mOsm at a pH of 7.3]. For voltage clamp recordings, neurons were voltage clamped at -70 mV using a Multiclamp 700B and currents were digitized with an Axon 1550B digitizer (Molecular Devices, Fremont, CA). For hM4d experiments, we included tetrodotoxin (500 nM), KA, picrotoxin (25 µM), and kynurenic acid (3 mM) in the bath to isolate the cell autonomous effects of the hM4d.

#### 4.5.3 Data analysis

Data were analyzed in Clampfit 11.1 (Molecular Devices, San Jose, CA). Membrane capacitance and resistance were determined online using a -10 mV square pulse after the cell stabilized (∼1 min). We did not correct for liquid junction potential.

### 4.6 Fiber Photometry Recordings

#### 4.6.1 Freely Behaving Recordings

In preparation for fiber photometry recordings in combination with drug treatments, mice were habituated to the behavior room, handling, exposure to i.p. injections, and patch cable attachment to their implanted fiber optics. In addition, mice individually took turns acclimating to the behavior chamber where photometry recordings take place. On experimental days, mice were brought into the behavior room and left undisturbed for 1-hour prior to recordings. A patch cord (200μm diameter, 0.37 NA; Newdoon; Shanghai, China) was then attached to the photometry system (FP3002, Neurophotometrics (MBF Biosciences); Williston, VT) and to mice’s fiber implants using a ceramic mating sleeve before placing them into the behavior chamber. A 415-nm LED (50 µW) was used to obtain isosbestic signal and 470-nm (50 µW) used to excite GCaMP7f where each wavelengths values were transcribed by Bonsai (Bonsai Foundation CIC, https://bonsai-rx.org). Alternating 415 and 470 nM LED stimulation was administered through the patch cable at 40 Hz. Each trial was 3 hours long where the 1st treatment was administered, i.p., at hour 1, the 2nd treatment administered at hour 2, followed by no more intervention and the recordings ending at hour 3. Treatment regimens were planned under a Latin-square design in order to prevent drug order effects. Each trial was separated at least 2 days to allow for drug wash-out.

#### 4.6.2 Fiber Photometry Data analysis

Fiber photometry data were analyzed using custom python notebooks and functions. Briefly, raw photometry data were deinterleaved to isolate 415 and 470 nM signals. Raw data were then visually inspected to identify major aberrations or signal disruptions due to patch cable hairpins or loss of signal due to optic fiber cannulae separation. Raw 415 and 470 nM signals were then independently fit using a biexponential function to correct for photobleaching of signal intensity. Fitted 415 signals were then regressed from the fitted 470 nm signals using Huber regression to remove motion-related isosbestic signals from the Ca-dependent signal. Processed 470 signals were then used for downstream alignment and normalization. We aligned user-generated timestamps for each injection to the processed 470 nm signal using a nearest-neighbor search function. Aligned timestamps are depicted as time point 0. We then selected 30 minute ranges prior to and after time point 0 and calculated raw and z-scored delta F/F. We examined the change of Z-Scored fluorescent values, looking at 30min pre-/ post-injection of the 2nd treatment administration to examine what effect pretreatment of the 1st treatment had on these values. For Net AUC analysis, we examined the change of Z-Score fluorescent values 15-30min post 2nd treatment administration for each individual animal, where the animal’s response to injection has subsided, using Kruskal-Wallis test with Dunn’s multiple comparisons. For Z-Scored peak response, we found the max Z-Scored value 0-30min post 2nd treatment injection for each individual animal and performed Kruskal-Wallis test with Dunn’s multiple comparisons. Python notebooks are available at https://github.com/Hardaway-Lab/FED3-Photometry_Workflows

### 4.7 Feeding Behavioral Assays

#### 4.7.1 FED3 Habituation and Fixed-Ratio 1 Training

Prior to feeding assays involving Feeding Experimental Devices (FED3), each mouse was singly housed in OneCages (Plexx; Netherlands) with their own FED3, where standard chow pellets were removed and instead, they fed on 20 mg precision grain pellets (Lab Diet; Cat no. 1811143 5TUM) [92]. In order to receive pellets, mice learned to operate the FED3 under a Fixed-Ratio 1 (FR1) paradigm, where they must nose-poke an active port on the FED3 to dispense an individual pellet. FED3s emit an auditory tone and deliver an LED light stimulus whenever the active port is selected to reinforce training. Mice were housed with their own FED3 and trained at least 1-week prior to any feeding assay. Mice were then tested following a fasted condition where they needed to consume at least 0.2 grams (10 precision pellets) within the 1st hour to ensure they are trained for refeeding experiments. Mice that failed to learn to feed from their FED3 or displayed or displayed food-hoarding behaviors were excluded from our studies. Unless otherwise specified, FED3s remained with the mice and fed ad libitum under FR1 paradigm for the remainder of our studies.

#### 4.7.2 Refeed

For food-deprived refeeding combined with drug treatments, mice were food-deprived for 24 hours. One hour before the 24-hour fasted time point, mice were weighed and assigned their randomized treatment regimen. Depending on the experiment, either one or two i.p. injections were administered prior to FED access. In two-treatment experiments, the first injection was given 40 minutes prior to FED3 access, followed by the second injection twenty minutes later. In single-treatment experiments, only one injection was given 20 minutes prior to FED3 access. Finally, at the 24-hour fasted time point, access to their FED was allowed where pellet retrieval and port selection was recorded by their FED3. After the 1st hour, pellets retrieved and consumed was visually confirmed by the experimenter. For two treatment experiments, regimens were planned under a Latin-square design in order to prevent drug order effects, with the exception of saline/saline administration which was performed first and served as a baseline for each mouse. Each session was separated at least 2 days to allow for drug wash-out.

#### 4.7.3 FED3 Data analysis

Data obtained from FR1 refeed studies were pulled from SD cards from each animal’s assigned FED3. The csv files obtained were then processed through FEDViz to bin their pellets retrieved and active pokes as a 1-hour timepoint [92]. These values were then recorded into spreadsheets for each treatment regime they received. Normalized values for pellets retrieved and active pokes were calculated individually for each animal by subtracting the values recorded at the 1-hour timepoint for each treatment regime it received, respectively, from the value recorded at their Veh/Veh treatment and then dividing it by their Veh/Veh value. These data were then converted into percentage and recorded into a spreadsheet as well along with their raw values. Raw values were left as is. Both normalized and raw values were then inputted into GraphPad Prism and analyzed using repeated measures two-way repeated measures ANOVA with Tukey’s multiple comparisons with the mean for each treatment compared to each other within their respective group as the multiple comparisons for both pellets retrieved and active pokes. When performing this analysis for normalized data sets, Veh/Veh values were set as zero and not included in the graphical plot.

#### 4.7.4 Intermittent HFD assay

To assess the consumption of palatable, calorie-dense food in the presence of neutral food, we generated a separate cohort of mice. Mice were singly housed 3 weeks after surgeries and kept on standard chow diet, ad libitum, throughout the study. Prior to the week of assays, crumbs of 60 kcal% fat rodent diet (HFD; Research Diets, Cat No. D12492,) were sprinkled into their cages to prevent neophagia on test days. On experimental days, mice were weighed and randomly assigned their treatment regimen. Mice were then injected, i.p., with their randomly assigned treatments before pre-weighed HFD and standard chow pellets were given 40 minutes later. We measured HFD and standard chow pellets weights an hour after HFD presentation and manually recorded the values for each mouse. Each session was separated at least 3 days to allow for drug wash-out and to avoid entrainment of HFD consumption.

#### 4.7.3 Intermittent HFD Data analysis

Data of HFD consumed for each individual mouse during intermittent HFD assays were recorded into spreadsheets to calculate the amount gram amount consumed, their normalized values, and the percentage of standard chow consumed. Normalized values for HFD consumption were calculated individually for each animal by subtracting the values recorded at the 1-hour timepoint for each treatment regime it received, respectively, from the value recorded at their Veh/Veh treatment and then dividing it by their Veh/Veh value. Similarly, percentage of standard chow consumed was calculated by dividing the gram amount of standard chow consumed with the gram amount of standard chow consumed plus gram amount of HFD consumed. These data were then converted into percentage and recorded into a spreadsheet as well along with their raw values. Raw values were left as is. Values were then inputted into Graph-Pad Prism and analyzed using repeated measures two-way repeated measures ANOVA with Tukey’s multiple comparisons with the mean for each treatment compared to each other within their respective group as the multiple comparisons. When performing this analysis for normalized data sets, Veh/Veh values were set as zero and not included in the graphical plot.

### 4.8 Imaging

Imaging was performed on a Keyence BZ-X810 under 20X magnification. For tissue sections obtained from C57BL6/J mice, tiled z-stacks of the CeA and surrounding region were captured using optical sectioning. For tissue sections obtained from any Cre mice, tiled z-stack images of the CeA and surrounding regions were captured using widefield fluorescence imaging. Raw images were then stitched and a maximum intensity projection made in Keyence Analyzer. Stitched raw images were obtained for individual channels including DAPI and 550 nm. Final images were assembled in Adobe Illustrator 25.2.3.

## Key Resource Table

| Reagent or Resource | Source | Identifier |
| --- | --- | --- |
| *Antibodies* |  |  |
| Rabbit anti-mcherry | Invitrogen, Thermo Fisher Scientific | Cat# PA5-34974; RRID:AB_2552323 |
| Cy5 AffiniPure Donkey anti-Rabbit | Jackson ImmunoResearch Labs | Cat# 711-175-152; RRID:AB_2340607 |
| Chicken anti-GFP | Aves Labs | Cat# GFP-1010; RRID:AB_2307313 |
| Alexa Fluor® 488 AffiniPure® Donkey Anti-Chicken IgY (IgG) (H+L) | Jackson ImmunoResearch Labs | Cat# PA5-34974; RRID:AB_2552323 |
| *Viral Strains* |  |  |
| pAAV-hSyn-DIO-mCherry (6.25*10<sup>12</sup> GC/ml) | Addgene | Cat# 50459-AAV9; RRID:Addgene_50459 |
| pAAV-hSyn-DIO-hM4D(Gi)-mCherry (5.75*10<sup>12</sup> GC/ml) | Addgene | Cat# 44362-AAV9; RRID:Addgene_44362 |
| pGP-AAV-syn-FLEX-jGCaMP7f-WPRE (2.8*10<sup>13</sup> GC/ml) | Addgene | Cat# 104492-AAV9; RRID:Addgene_104492 |
| *Chemicals, peptides, and recombinant proteins* |  |  |
| Deschloroclozapine dihydrochloride | HelloBio | Cat# HB9126 |
| Exendin-4 | MedChemExpress | Cat# HY-13443 |
| Exendin(9-39) amide | MedChemExpress | Cat# HY-P0264 |
| Normal Donkey Serum | Jackson ImmunoResearch Labs | Cat# 017-000-121; RRID: AB_2337258 |
| Bovine Serum Albumin | Sigma-Aldrich | Cat# A2153 |
| VECTASHIELD® HardSet™ Antifade Mounting Medium with DAPI | Vector Laboratories | Cat# H-1500-10 |
| Triton X-100 | Fisher Scientific | Cat# BP151-500 |
| Kynurenic acid | MedChemExpress | Cat# HY-100806 |
| *Experimental models: Organisms/strains* |  |  |
| Mouse: B6J.129S6(FVB)-Slc32a1<sup>tm2(cre)Lowl/MwarJ</sup> | The Jackson Laboratory | RRID:IMSR_JAX:028862 |
| Mouse: STOCK Tg(Prkcd-glc-1/CF-P,-cre)EH124Gsat/Mmucd | MMRRC | RRID:MMRRC_011559-UCD |
| Mouse: STOCK Ssttm2.1(cre)Zjh/J | The Jackson Laboratory | RRID:IMSR_JAX:013044 |
| Mouse: STOCK Glp1rtm1.1(cre)Lbrl/RcngJ | The Jackson Laboratory | RRID:IMSR_JAX:029283 |
| *Software and algorithms* |  |  |
| Graphpad Prism v10 | GraphPad | https://www.graphpad.com/; RRID:SCR_002798 |
| ImageJ/FIJI | NIH | https://fiji.sc/; RRID:SCR_002285 |
| Adobe Illustrator | Adobe | https://www.adobe.com/products/illustrator.html; RRID:SCR_010279 |
| pCLAMP v10 | Molecular Devices | http://www.moleculardevices.com/products/software/pclamp.html; RRID:SCR_011323 |
| Microsoft Excel | Microsoft | www.microsoft.com/en-gb/; RRID:SCR_016137 |
| FED3_Viz | Lex Kravitz Lab | https://github.com/earnestt1234/FED3_Viz |
| Bonsai | Bonsai Foundation CIC | https://bonsai-rx.org/; RRID:SCR_017218 |
| *Other* |  |  |
| MLABRODENT TABLET 20mg, Grain-Based Rodent Tablet | TestDiet, Lab Supply Orders | CAT# 1811143 5TUM |
| Fiber Optic Cannula | Newdoon | Cat# FOC-C-B-200-1.25-0.37-10 |
| Rodent Diet With 60 kcal% Fat, D12492 | Research Diets Inc | Cat# D12492 |
| C&B Metabond® Quick Adhesive Cement System | Parkell | Cat# S380 |
| Dental Cement | Stoelting Co. |  |
| Nanoject III | Drummond Scientific Company | Cat# 3-000-207 |
| Nanoject Glass Capillaries | Drummond Scientific Company | Cat# 3-000-203-G/X |
| Screws | McMaster-Carr | Cat# CS-335 |
| Fiber optic cannulae holder | This paper | https://github.com/OpenBehavior/Modular-Stereotaxic-Holders |
| Feeding Experimental Device 3 | Handmade or OpenEphys | https://open-ephys.org/ |
| Fiber-optic Patch Cords | Doric Lenses | Cat# BB-P(2)_200/220/900/900-0.37_3m_FCM-2xMF1.25(F) |
| FP3002 | Neurophotometrics, MBF Bioscience | https://neurophotometrics.com/fp3002 |

---

**Supplement Figure 1.**
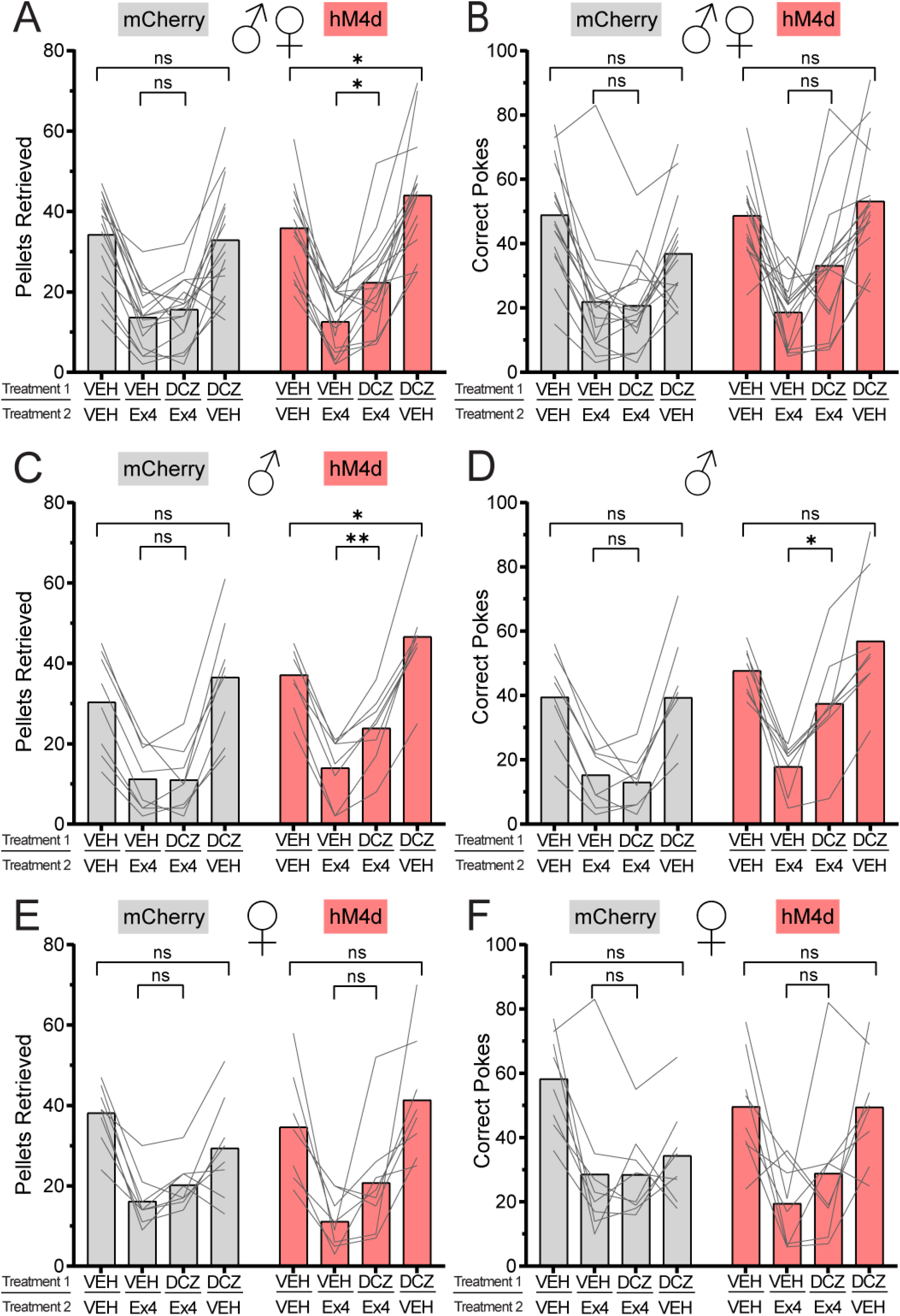
Chemogenetic inhibition of CeA neurons attenuates systemic Exendin-4-mediated hypophagia. **A**. Pellets retrieved by all *vGat*-Cre mice during each treatment session. Group (mCherry/hM4d) X Treatment: F(3, 90) = 3.772, *p* = 0.0134; Group effect: F(1, 30) = 2.450, *p* = 0.1280; Treatment effect: F(2.452, 73.56) = 77.75, *p* < 0.0001. Post tests - Veh/Ex4 vs DCZ/Ex4: mCherry (*p* = 0.5348), hM4d (*p* = 0.0157); DCZ/Veh vs Veh/Veh: mCherry (*p* = 0.9756), hM4d (*p* = 0.0257). **B**. Active port pokes by all *vGat*-Cre mice during each treatment session. Group (mCherry/hM4d) X Treatment: F (3, 90) = 4.574, *p* = 0.0050; Group effect: F (1, 30) = 2.258, *p* = 0.1433; Treatment effect: F (2.880, 86.40) = 38.49, *p* < 0.0001. Post tests - Veh/Ex4 vs DCZ/Ex4: mCherry (*p* = 0.9791), hM4d (*p* = 0.0553); DCZ/Veh vs Veh/Veh: mCherry (*p* = 0.1123), hM4d (*p* = 0.7854). **C**. Pellets retrieved by male *vGat*-Cre mice during each treatment session. Group (mCherry/hM4d) X Treatment: F(3, 42) = 2.162, *p* = 0.1068; Group effect: F(1, 14) = 3.201, *p* = 0.0952; Treatment effect: F(3, 42) = 83.55, *p* < 0.0001. Post tests - Veh/Ex4 vs DCZ/Ex4: mCherry (*p* = 0.9998), hM4d (*p* = 0.0097); DCZ/Veh vs Veh/Veh: mCherry (*p* = 0.1830), hM4d (*p* = 0.0136). **D**. Active port pokes by male *vGat*-Cre mice during each treatment session. Group (mCherry/hM4d) X Treatment: F(3, 42) = 4.027, *p* = 0.0132; Group effect: F(1, 14) = 6.356, *p* = 0.0244; Treatment effect: F(2.294, 32.12) = 38.27, *p* < 0.0001. Post tests: Veh/Ex4 vs DCZ/Ex4: mCherry (*p* = 0.8596), hM4d (*p* = 0.0235); DCZ/Veh vs Veh/Veh: mCherry (p > 0.9999), hM4d (*p* = 0.6571). **E**. Pellets retrieved by female *vGat*-Cre mice during each treatment session. Group (mCherry/hM4d) X Treatment: F(2.349, 32.89) = 2.910, *p* = 0.0608; Group effect: F(1, 14) = 0.07153, *p* = 0.7930; Treatment effect: F (2.349, 32.89) = 24.79, *p* < 0.0001. Post tests: Veh/Ex4 vs DCZ/Ex4: mCherry (*p* = 0.1511), hM4d (*p* = 0.3689); DCZ/Veh vs Veh/Veh: mCherry (*p* = 0.2489), hM4d (*p* = 0.1096). **F**. Active port pokes by female *vGat*-Cre mice during each treatment session. Group (mCherry/hM4d) X Treatment: F(3, 42) = 2.523, *p* = 0.0706; Group effect: F(1, 14) = 0.007972, *p* = 0.9301; Treatment effect: F(2.482, 34.75) = 14.35, *p* < 0.0001. Post tests: Veh/Ex4 vs DCZ/Ex4: mCherry (*p* >0.9999), hM4d (*p* = 0.7288); DCZ/Veh vs Veh/Veh: mCherry (*p* = 0.0569), hM4d (*p* > 0.9999). For **A-F**, data were analyzed using two-way repeated measures ANOVA with Tukey’s multiple comparisons.

**Supplement Figure 2.**
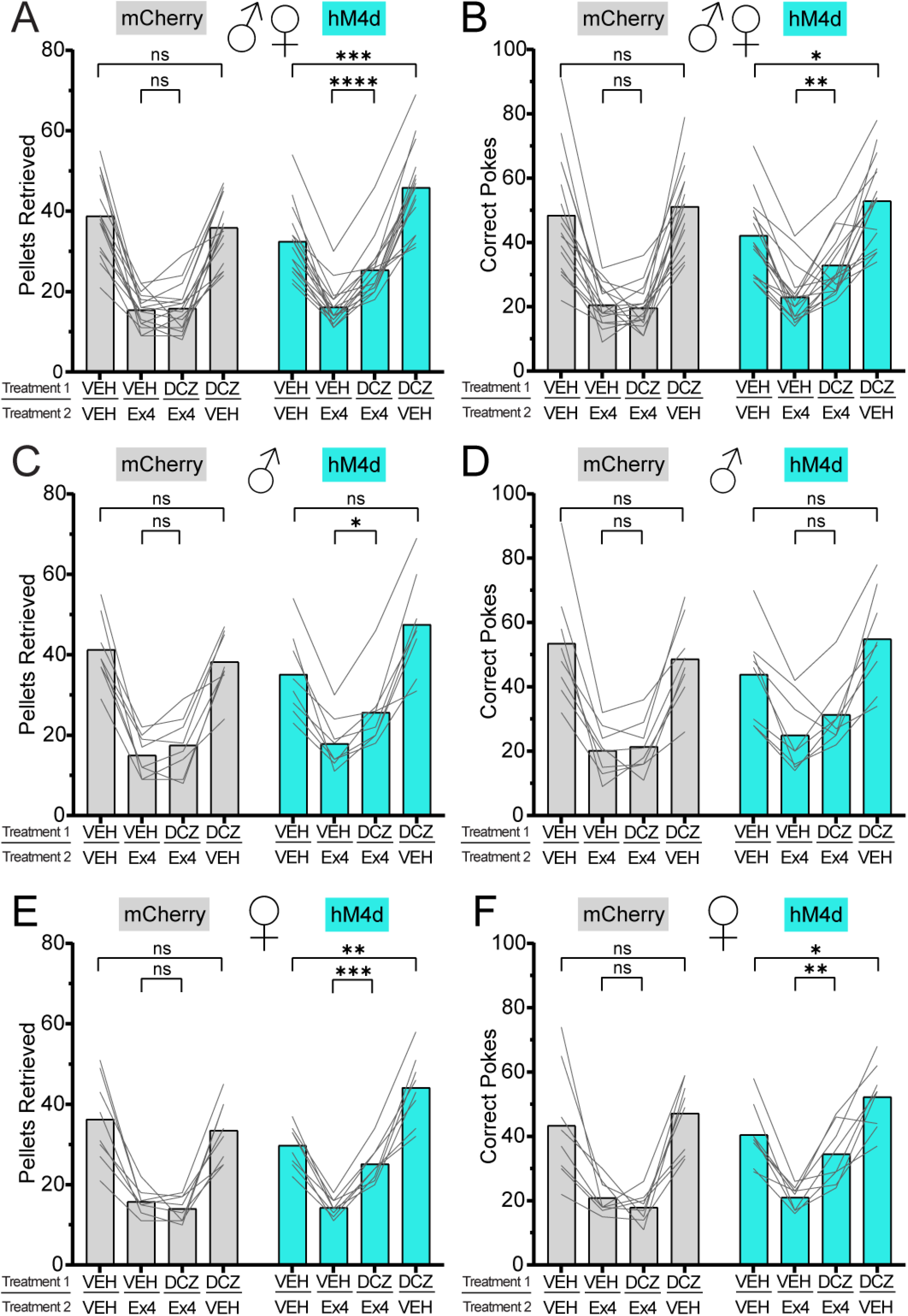
Chemogenetic inhibition of *Prkcd*^CeA^ neurons potently attenuates systemic Exendin-4-mediated hypophagia. **A**. Pellets retrieved by all *Prkcd*-Cre mice during each treatment session. Group (mCherry/hM4d) X Treatment: F(3, 90) = 15.59, *p* < 0.0001; Group effect: F(1, 30) = 2.882, *p* = 0.0999; Treatment effect: F(2.324, 69.73) = 145.9, *p* < 0.0001. Post tests: Veh/Ex4 vs DCZ/Ex4: mCherry (*p* = 0.9871), hM4d (*p* < 0.0001); DCZ/Veh vs Veh/Veh: mCherry (*p* = 0.3863), hM4d (*p* = 0.0002) **B**. Active port pokes by all *Prkcd*-Cre mice during each treatment session. Group (mCherry/hM4d) X Treatment: F(3, 90) = 5.840, *p* = 0.0011; Group effect: F(1, 30) = 0.9368, *p* = 0.3408; Treatment effect: F(2.512, 75.36) = 77.30, *p* < 0.0001. Post tests: Veh/Ex4 vs DCZ/Ex4: mCherry (*p* = 0.9550), hM4d (*p* = 0.0077); DCZ/Veh vs Veh/Veh: mCherry (*p* = 0.9418), hM4d (*p* = 0.0150) **C**. Pellets retrieved by male *Prkcd*-Cre mice during each treatment session. Group (mCherry/hM4d) X Treatment: F(3, 42) = 5.213, *p* = 0.0038; Group effect: F(1, 14) = 1.091, *p* = 0.3139; Treatment effect: F(2.325, 32.55) = 68.95, *p* < 0.0001. Post tests: Veh/Ex4 vs DCZ/Ex4: mCherry (*p* = 0.3804), hM4d (*p* = 0.0126); DCZ/Veh vs Veh/Veh: mCherry (*p* = 0.6885), hM4d (*p* = 0.0835) **D**. Active port pokes by male *Prkcd*-Cre mice during each treatment session. Group (mCherry/hM4d) X Treatment: F(3, 42) = 3.361, *p* = 0.0275; Group effect: F(1, 14) = 0.3191, *p* = 0.5811; Treatment effect: F(2.218, 31.06) = 38.10, *p* < 0.0001. Post tests: Veh/Ex4 vs DCZ/Ex4: mCherry (*p* = 0.9992), hM4d (*p* = 0.3458); DCZ/Veh vs Veh/Veh: mCherry (*p* = 0.8301), hM4d (*p* = 0.5580) **E**. Pellets retrieved by female *Prkcd*-Cre mice during each treatment session. Group (mCherry/hM4d) X Treatment: F(3, 42) = 11.89, *p* < 0.0001; Group effect: F(1, 14) = 2.277, *p* = 0.1535; Treatment effect: F(2.043, 28.60) = 76.10, *p* < 0.0001. Post tests: Veh/Ex4 vs DCZ/Ex4: mCherry (*p* = 0.6469), hM4d (*p* = 0.0003); DCZ/Veh vs Veh/Veh: mCherry (*p* = 0.6879), hM4d (*p* = 0.0013). **F**. Active port pokes by female *Prkcd*-Cre mice during each treatment session. Group (mCherry/hM4d) X Treatment: F(3, 45) = 4.874, *p* = 0.0051; Group effect: F(1, 15) = 3.074, *p* = 0.1000; Treatment effect: F(2.218, 33.27) = 43.40, *p* < 0.0001. Post tests: Veh/Ex4 vs DCZ/Ex4: mCherry (*p* = 0.7466), hM4d (*p* = 0.0097); DCZ/Veh vs Veh/Veh: mCherry (*p* = 0.8903), hM4d (*p* = 0.0183). For **A-F**, data were analyzed using two-way repeated measures ANOVA with Tukey’s multiple comparisons.

**Supplement Figure 3.**
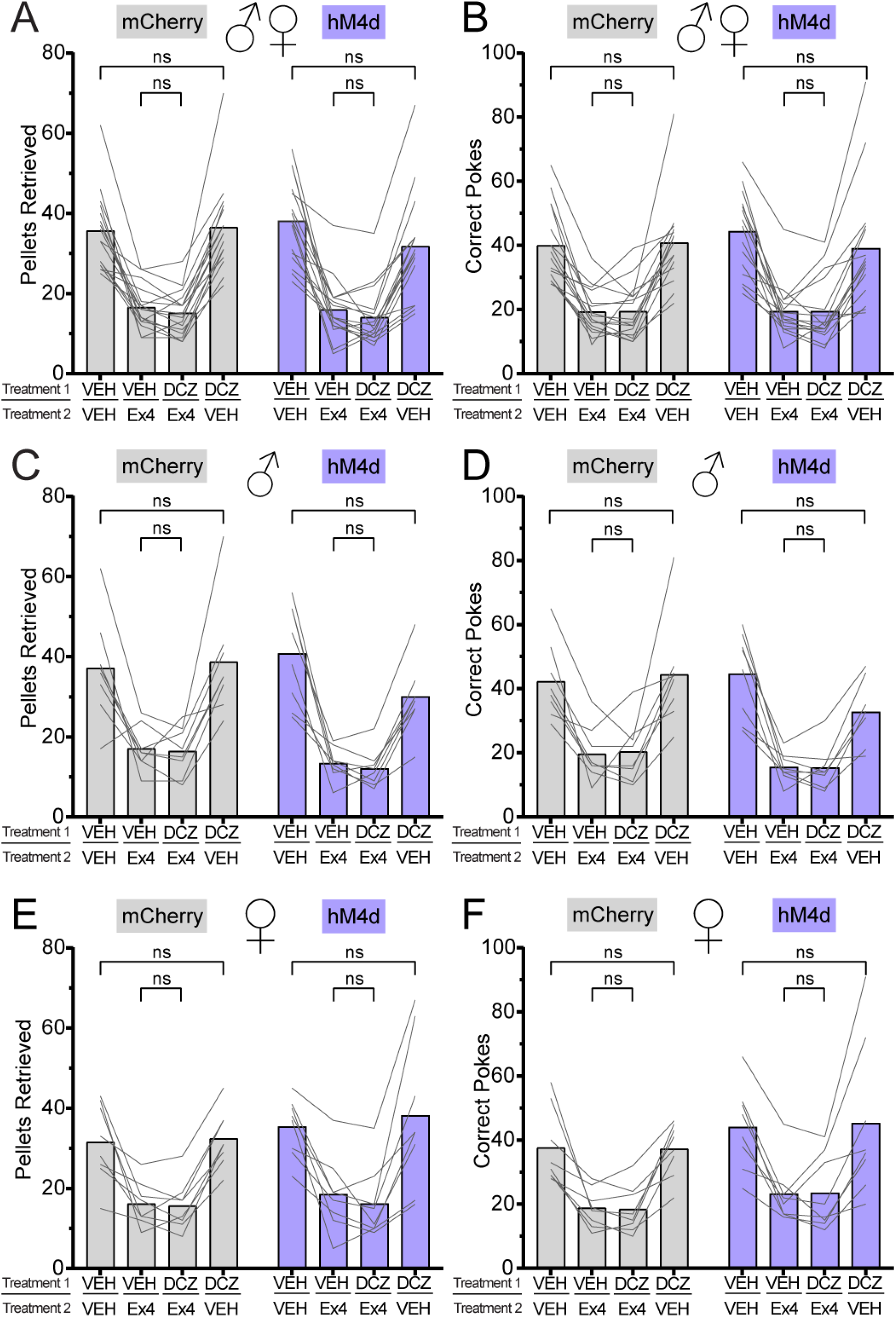
Chemogenetic inhibition of *Sst*^CeA^ neurons has no effect on systemic Exendin-4-mediated hypophagia. **A**. Pellets retrieved by all *Sst*-Cre mice during each treatment session. Group (mCherry/hM4d) X Treatment: F(2.124, 63.72) = 1.451, *p* = 0.2415; Group effect: F(1, 30) = 0.1609, *p* = 0.6912; Treatment effect: F(2.124, 63.72) = 90.86, *p* < 0.0001. Post tests: Veh/Ex4 vs DCZ/Ex4: mCherry (*p* = 0.4255), hM4d (*p* = 0.4421); DCZ/Veh vs Veh/Veh: mCherry (*p* = 0.9739), hM4d (*p* = 0.3669). **B**. Active port pokes by all *Sst*-Cre mice during each treatment session. Group (mCherry/hM4d) X Treatment: F(2.451, 73.52) = 0.7707, *p* = 0.4904; Group effect: F(1, 30) = 0.04180, *p* = 0.8394; Treatment effect: F(2.451, 73.52) = 70.53, *p* < 0.0001,. Post tests: Veh/Ex4 vs DCZ/Ex4: mCherry (*p* = 0.9999), hM4d (*p* > 0.9999); DCZ/Veh vs Veh/Veh: mCherry (*p* = 0.9864), hM4d (*p* = 0.5825). **C**. Pellets retrieved by male *Sst*-Cre mice during each treatment session. Group (mCherry/hM4d) X Treatment: F(1.989, 27.85) = 1.784, *p* = 0.1867; Group effect: F(1, 14) = 0.9185, *p* = 0.3541; Treatment effect: F(1.989, 27.85) = 45.32, *p* < 0.0001. Post tests: Veh/Ex4 vs DCZ/Ex4: mCherry (*p* > 0.9999), hM4d (*p* > 0.9999); DCZ/Veh vs Veh/Veh: mCherry (*p* > 0.9999), hM4d (*p* = 0.2347). **D**. Active port pokes by male *Sst*-Cre mice during each treatment session. Group (mCherry/hM4d) X Treatment: F(2.239, 31.34) = 1.928, *p* = 0.1581; Group effect: F(1, 14) = 1.349, *p* = 0.2649 Treatment effect: F(2.239, 31.34) = 43.49, *p* < 0.0001. Post tests: Veh/Ex4 vs DCZ/Ex4: mCherry (*p* = 0.9956), hM4d (*p* = 0.9990); DCZ/Veh vs Veh/Veh: mCherry (*p* = 0.9443), hM4d (*p* = 0.0728). **E**. Pellets retrieved by female *Sst*-Cre mice during each treatment session. Group (mCherry/hM4d) X Treatment: F(1.763, 24.68) = 0.3736, *p* = 0.6659; Group effect: F(1, 14) = 0.6553, *p* = 0.4318; Treatment effect: F(1.763, 24.68) = 31.86, *p* < 0.0001. Post tests: Veh/Ex4 vs DCZ/Ex4: mCherry (*p* = 0.9844), hM4d (*p* = 0.6667); DCZ/Veh vs Veh/Veh: mCherry (*p* = 0.9731), hM4d (*p* = 0.6667). **F**. Active port pokes by female *Sst*-Cre mice during each treatment session. Group (mCherry/hM4d) X Treatment: F (2.280, 31.92) = 0.1359, *p* = 0.8970; Group effect: F(1, 14) = 1.369, *p* = 0.2615; Treatment effect: F(2.280, 31.92) = 28.93, *p* < 0.0001. Post tests: Veh/Ex4 vs DCZ/Ex4: mCherry (*p* = 0.9823), hM4d (*p* = 0.9998); DCZ/Veh vs Veh/Veh: mCherry (*p* = 0.9996), hM4d (*p* = 0.9975). For **A-F**, data were analyzed using two-way repeated measures ANOVA with Tukey’s multiple comparisons.

**Supplement Figure 4.**
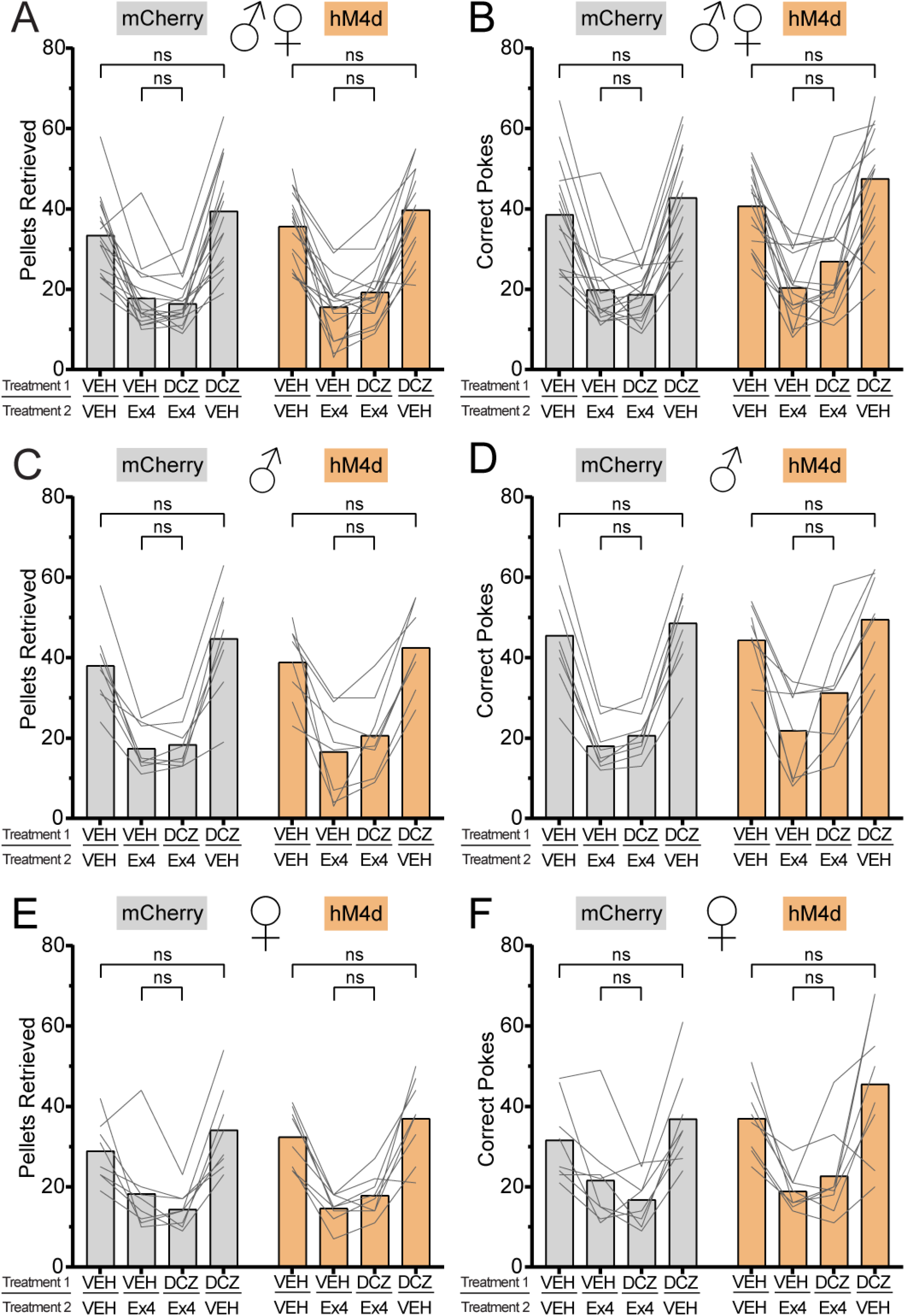
Chemogenetic inhibition of *Glp1r*^CeA^ neurons attenuates systemic Exendin-4-mediated hypophagia. **A**. Pellets retrieved by all *Glp1r*-Cre mice during each treatment session. Group (mCherry/hM4d) X Treatment: F(2.189, 65.68) = 0.9496, *p* = 0.3991; Group effect: F(1, 30) = 0.09712, *p* = 0.7575; Treatment effect: F(2.189, 65.68) = 100.0, *p* < 0.0001. Post tests: Veh/Ex4 vs DCZ/Ex4: mCherry (*p* = 0.7897), hM4d (*p* = 0.1046); DCZ/Veh vs Veh/Veh: mCherry (*p* = 0.0679), hM4d (*p* = 0.2579). **B**. Active port pokes by all *Glp1r*-Cre mice selected during each treatment session. Group (mCherry/hM4d) X Treatment: F(2.684, 80.51) = 1.256, *p* = 0.2947; Group effect: F(1, 30) = 1.672, *p* = 0.2059; Treatment effect: F(2.684, 80.51) = 67.57, *p* < 0.0001. Post tests: Veh/Ex4 vs DCZ/Ex4: mCherry (*p* = 0.9451), hM4d (*p* = 0.1496); DCZ/Veh vs Veh/Veh: mCherry (*p* = 0.3263), hM4d (*p* = 0.2298). **C**. Pellets retrieved by male *Glp1r*-Cre mice during each treatment session. Group (mCherry/hM4d) X Treatment: F(1.963, 27.49) = 0.3547, *p* = 0.7007; Group effect: F(1, 14) = 6.147e-005, *p* = 0.9939; Treatment effect: F(1.963, 27.49) = 66.38, *p* < 0.0001. Post tests: Veh/Ex4 vs DCZ/Ex4: mCherry (*p* = 0.7722), hM4d (*p* = 0.4944); DCZ/Veh vs Veh/Veh: mCherry (*p* = 0.1473), hM4d (*p* = 0.6130). **D**. Active port pokes by male *Glp1r*-Cre mice during each treatment session. Group (mCherry/hM4d) X Treatment: F(2.630, 36.83) = 2.221, *p* = 0.1092; Group effect: F(1, 14) = 0.6753, *p* = 0.4250; Treatment effect: F(2.630, 36.83) = 67.95, *p* < 0.0001. Post tests: Veh/Ex4 vs DCZ/Ex4: mCherry (*p* = 0.0853), hM4d (*p* = 0.2834); DCZ/Veh vs Veh/Veh: mCherry (*p* = 0.8054), hM4d (*p* = 0.4777). **E**. Pellets retrieved by female *Glp1r*-Cre mice during each treatment session. Group (mCherry/hM4d) X Treatment: F(2.475, 34.64) = 1.191, *p* = 0.3227; Group effect: F(1, 14) = 0.2856, *p* = 0.6014; Treatment effect: F(2.475, 34.64) = 38.88, *p* < 0.0001. Post tests: Veh/Ex4 vs DCZ/Ex4: mCherry (*p* = 0.5282), hM4d (*p* = 0.1664); DCZ/Veh vs Veh/Veh: mCherry (*p* = 0.5176), hM4d (*p* = 0.5482). **F**. Active port pokes by female *Glp1r*-Cre mice during each treatment session. Group (mCherry/hM4d) X Treatment: F(2.217, 31.03) = 1.109, *p* = 0.3477; Group effect: F(1, 14) = 1.199, *p* = 0.2920; Treatment effect: F(2.217, 31.03) = 20.93, *p* < 0.0001. Post tests: Veh/Ex4 vs DCZ/Ex4: mCherry (*p* = 0.5651), hM4d (*p* = 0.6817); DCZ/Veh vs Veh/Veh: mCherry (*p* = 0.4680), hM4d (*p* = 0.5409). For **A-F**, data were analyzed using two-way repeated measures ANOVA with Tukey’s multiple comparisons.

**Supplement Figure 5.**
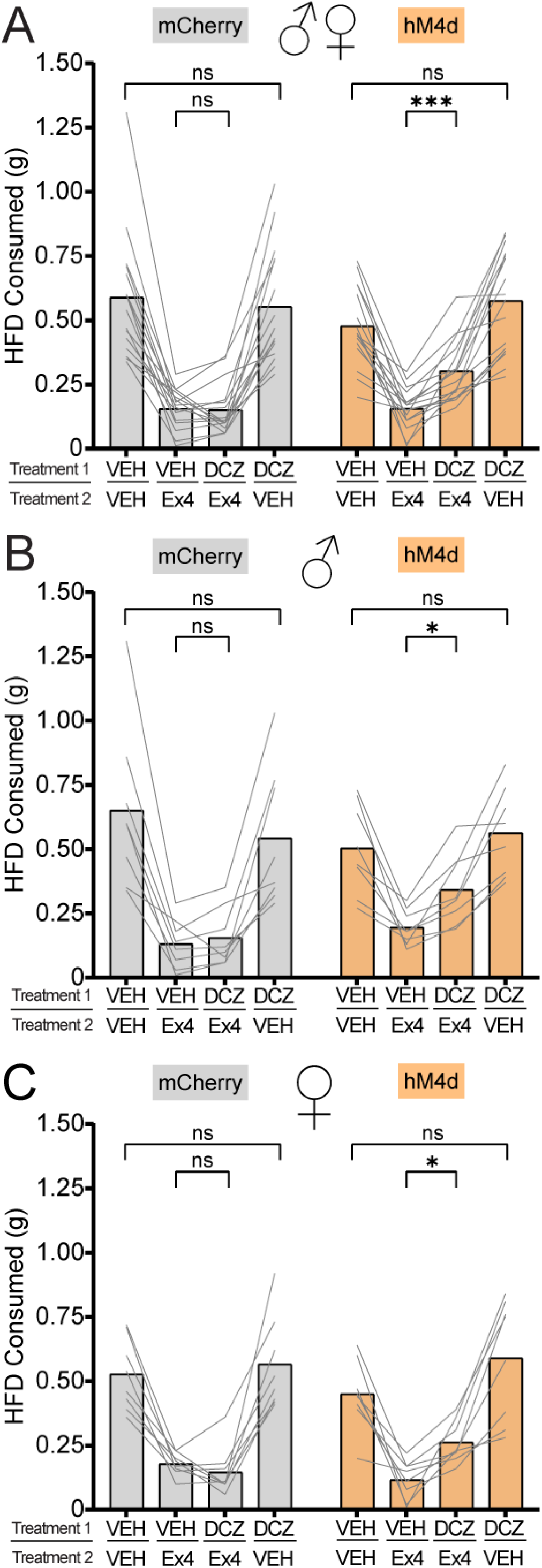
Chemogenetic inhibition of *Glp1r*^CeA^ neurons attenuates systemic Exendin-4-mediated reduction in palatable food intake. **A**. Grams of high-fat pellet consumed by all *Glp1r*-Cre mice during each treatment session. Group (mCherry/hM4d) X Treatment: F (2.303, 69.08) = 5.455, *p* = 0.0044; Group effect: F(1, 30) = 0.1463, *p* = 0.7048; Treatment effect: F(2.303, 69.08) = 81.92, *p* < 0.0001. Post test: Veh/Ex4 vs DCZ/Ex4: mCherry (*p* = 0.9979), hM4d (*p* = 0.0001); DCZ/Veh vs Veh/Veh: mCherry (*p* = 0.9293), hM4d (*p* = 0.2442). **B**. Grams of high-fat pellet consumed by *Glp1r*-Cre male mice during each treatment session. Group (mCherry/hM4d) X Treatment: F(2.237, 31.31) = 3.787, *p* = 0.0296; Group effect: F(1, 14) = 0.1921, *p* = 0.6678; Treatment effect: F(2.237, 31.31) = 35.25, *p* < 0.0001. Post test: Veh/Ex4 vs DCZ/Ex4: mCherry (*p* = 0.7702), hM4d (*p* = 0.0195); DCZ/Veh vs Veh/Veh: mCherry (*p* = 0.5843), hM4d (*p* = 0.8714). **C**. Grams of high-fat pellet consumed by *Glp1r*-Cre female mice during each treatment session. Group (mCherry/hM4d) X Treatment: F(2.197, 30.76) = 2.249, *p* = 0.1182; Group effect: F(1, 14) = 4.891e-005, *p* = 0.9945; Treatment effect: F(2.197, 30.76) = 50.17, *p* < 0.0001. Post test: Veh/Ex4 vs DCZ/Ex4: mCherry (*p* = 0.7405), hM4d (*p* = 0.0176); DCZ/Veh vs Veh/Veh: mCherry (*p* = 0.9562), hM4d (*p* = 0.2309). For **A-C**, data were analyzed using two-way repeated measures ANOVA with Tukey’s multiple comparisons.

